# SFPQ dysregulation promotes TDP-43 pathology through a pathogenic feedback loop

**DOI:** 10.64898/2026.07.22.740184

**Authors:** Alison L Hogan, Patrick Chiu, Natalie Grima, Sharlynn Wu, Grant Ritcher, Cindy Maurel, Madison Kane, Andrew Smith, Shu Yang, Adam K Walker, Ian P Blair, Marco Morsch

## Abstract

TDP-43 proteinopathies comprise a group of clinically distinct neurodegenerative disorders unified by common pathological changes in the RNA-binding protein TDP-43, including nuclear loss of function (LOF), cytoplasmic mislocalisation and aggregation. The presence of these hallmarks across diverse diseases, such as frontotemporal dementia (FTD), amyotrophic lateral sclerosis (ALS), alzheimer’s disease (AD) and Limbic-predominant Age-related TDP-43 Encephalopathy (LATE), suggests the involvement of convergent upstream regulatory mechanisms. This study identified SFPQ (Splicing Factor Proline and Glutamine Rich) as one such factor.

The depletion of nuclear SFPQ, in addition to its cytoplasmic accumulation and aggregation has been described across ALS, AD and FTD patient studies and in multiple genetic models of these diseases. Evidence to date places SFPQ pathology downstream of TDP-43 dysfunction. This study provided further evidence that TDP-43 drives SFPQ pathology while revealing a reciprocal role for SFPQ in regulating TDP-43 homeostasis. Our data shoed that SFPQ LOF was associated with a shift in TDP-43 RNA isoform usage away from the canonical protein-coding transcript towards isoforms predicted to undergo nonsense-mediated decay (NMD). Consistent with this, TDP-43 protein expression was reduced across multiple model systems in which SFPQ expression was suppressed. Furthermore, cytoplasmic accumulation of SFPQ was found to promote the mislocalisation of TDP-43 and its sequestration within SFPQ-containing cytoplasmic condensates that exhibited progressively reduced molecular mobility over time.

Collectively, these findings identify SFPQ as an active contributor to multiple facets of TDP-43 pathology, support a bidirectional relationship between the two proteins, and suggest the existence of a feed-forward, self-amplifying mechanism that may contribute to disease progression in TDP-43 proteinopathies.

**Graphical Abstract:** 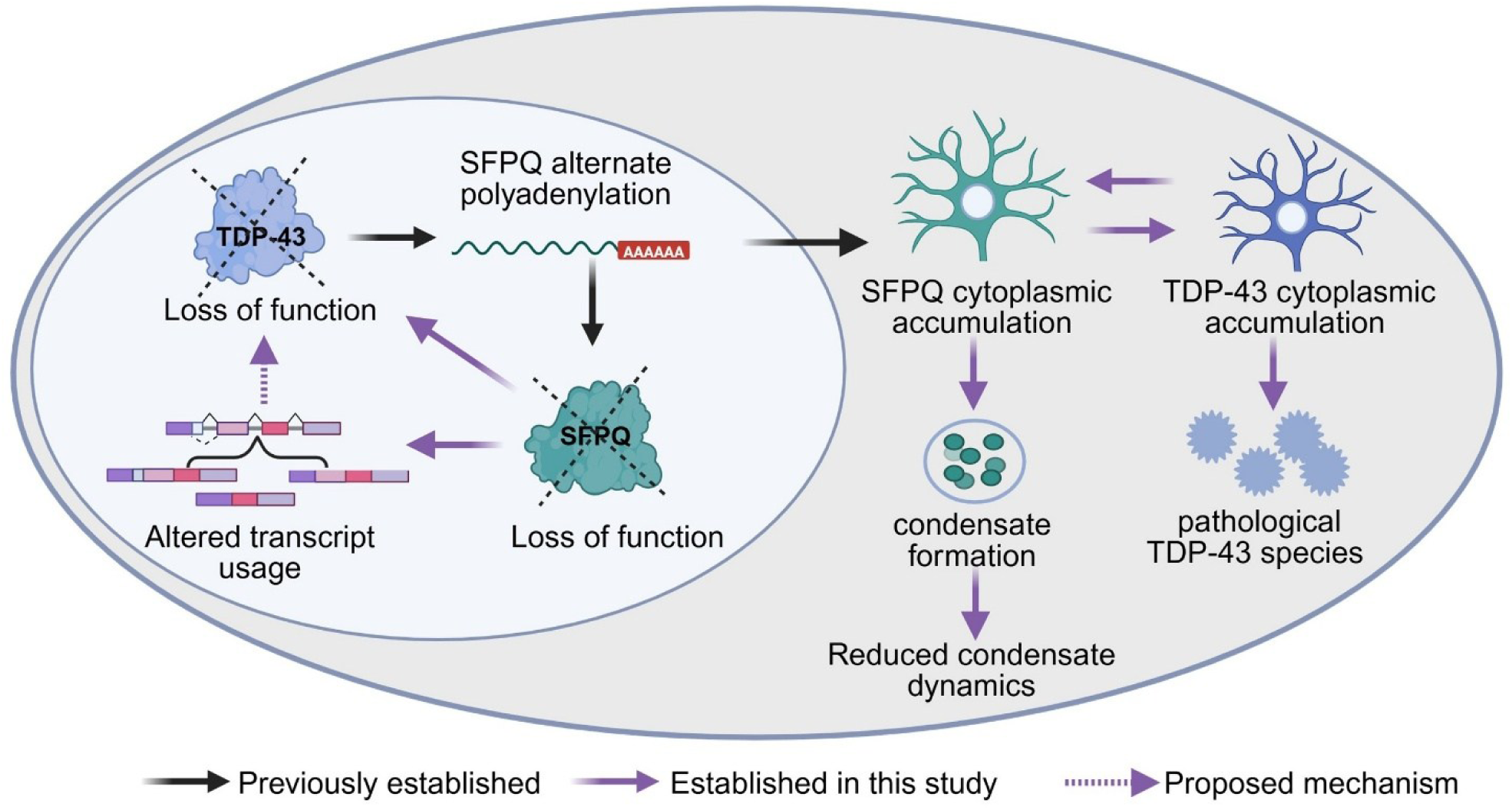

## Introduction

TAR DNA-binding protein 43 (TDP-43) is an DNA and RNA-binding protein that undergoes characteristic pathological changes across multiple neurodegenerative diseases including amyotrophic lateral sclerosis, ALS (>90% of patients^1^), frontotemporal lobar degeneration, FTLD (≤ 50% of patients^2^), Limbic-predominant Age-related TDP-43 Encephalopathy, LATE (100% of patients) and Alzheimers disease, AD (∼57% of patients^3^). These pathological changes include TDP-43 nuclear loss of function (LOF), cytoplasmic accumulation, post-translational modifications (including hyperphosphorylation) and aggregation into ubiquitinated, insoluble inclusions^4^.

The commonality of TDP-43 pathology across multiple disorders underscores its central role in neurodegeneration. Extensive work has therefore focused on defining the functional consequences of TDP-43 perturbation, revealing both gain-and loss-of-function (LOF) effects. Cytoplasmic accumulation of TDP-43 promotes the formation of insoluble aggregates that sequester functional proteins, disrupt nuclear–cytoplasmic transport, and activate cell death pathways^5^, Concurrently, nuclear TDP-43 LOF leads to widespread RNA processing defects, including cryptic exon retention^6–8^ and alternate polyadenylation^9^ and hyper-assembly of paraspeckles - nuclear ribonucleoprotein condensates that regulate gene expression through selective RNA and protein sequesteration^10^. While these downstream consequences of TDP-43 nuclear loss and mislocalisation are well characterised, the upstream factors that initiate and propagate these pathological changes remain incompletely defined.

Splicing Factor Proline-and Glutamine-Rich (SFPQ) is a predominantly nuclear DNA and RNA-binding protein with essential roles in transcriptional regulation, elongation of long genes, alternative splicing, RNA transport, DNA repair, and paraspeckle formation (reviewed in ^11^). SFPQ dysregulation has emerged as a recurring feature across diverse models of neurodegeneration: nuclear depletion of SFPQ has been reported in ALS models carrying mutations in *TARDBP* (M337V pig^12^), *SOD1 (*G93A mouse^13^), *VCP (*A232E mouse and iPSC-derived neurons^13^), and *CCNF* (S621G HEK-293T cells^14^), as well as in the Tau P301L mouse model of AD^15^. In human tissue, SFPQ nuclear loss^12,13^ and cytoplasmic aggregation^16^ have been reported in ALS-affected motor neurons, with nuclear depletion also reported across multiple dementias including AD and FTD^15,17,18^. Together, these findings suggest that disruption of SFPQ homeostasis is a common pathological feature across TDP-43 proteinopathies.

Mechanistic links between TDP-43 and SFPQ have begun to emerge^9,19^. *In vitro* studies have determined that nuclear TDP-43 regulates SFPQ polyadenylation by binding to its 3′ untranslated region (UTR), thereby promoting use of a proximal polyadenylation site. TDP-43 nuclear depletion causes a shift to a more distal polyadenylation site which generates transcripts that are targeted for degradation via nonsense-mediated decay (NMD).

Transcripts that escape this process are translated into a SFPQ isoform that lacks a nuclear localisation sequence (NLS) and consequently displays increased cytoplasmic expression. Notably, the alternately polyadenylated SFPQ transcript has been detected in human induced neurons and ALS patient frontal cortex^19^, confirming the clinical relevance of the SFPQ-TDP-43 relationship.

Here, we examine the relationship between TDP-43 pathology and SFPQ dysfunction in the well-characterised TDP-43^rNLS8^ mouse model^20^, demonstrating for the first time in a mammalian system that TDP-43 pathology is accompanied by SFPQ LOF and cytoplasmic mislocalisation. Additionally, using a combination of cell and zebrafish models, we establish that SFPQ is not merely a downstream consequence of TDP-43 dysregulation, but actively contributes to its progression. We demonstrate that SFPQ LOF alters *TARDBP* RNA processing and reduces TDP-43 protein expression. In parallel, we show that upon accumulation in the cytoplasm, SFPQ assembles into biomolecular condensates that can sequester TDP-43 and over time transition to a state of reduced solubility. Further, we demonstrate that cytoplasmic accumulation of SFPQ can drive mislocalisation of TDP-43 in motor neurons *in vivo* and over time leads to formation of pathological TDP-43 species in motor neurons. Collectively, these findings identify SFPQ as a driver of key pathological features of TDP-43 proteinopathies, supporting a bidirectional, feed-forward relationship between SFPQ and TDP-43 that may contribute to disease progression.

## Results

### TDP-43 pathology causes reduced expression and mislocalisation of SFPQ that precedes neurodegeneration

*In vitro* TDP-43 knockdown studies have identified a link between TDP-43 LOF and production of an alternately polyadenylated SFPQ transcript^9,19^. Notably, this transcript is a target of NMD, while transcripts that escape this process encode an SFPQ protein isoform lacking a NLS. Accordingly, alternative polyadenylation contributes to both SFPQ LOF and its cytoplasmic accumulation. While these findings provide a mechanistic framework linking TDP-43 dysfunction to SFPQ pathology, this relationship has yet to be demonstrated *in vivo*, and the temporal sequence of TDP-43–driven SFPQ pathology remains undefined. To address this, we utilised an inducible model expressing TDP-43 with a mutated NLS, the TDP-43^rNLS8^ mouse^20^.

The TDP-43^rNLS^^8^ mouse exhibits both cytoplasmic accumulation and approximately 50% nuclear depletion of TDP-43, while developing a rapid and severe phenotype in which neuronal death is reported to become evident at 6 weeks post induction^20^. To examine progressive changes to SFPQ in this model, spinal cord tissue was collected at 2 weeks post induction (representing disease onset), 4 weeks post induction, and 6 weeks post induction.

We first confirmed that neuronal death was evident in our mouse cohort at the reported 6 week timepoint (Supplementary Fig.1A). We then quantified SFPQ neuronal expression and localisation using immunofluorescence (**Fig. 1A**). SFPQ whole cell expression significantly reduced progressively from the 2-week timepoint (mean = 7150 ± 622, n = 4 mice, n > 56 neurons per mouse) to the 4-week timepoint (mean = 5094 ± 1139, n = 6 mice, n > 121 neurons per mouse, p <0.0001), and was further reduced at the 6-week timepoint (mean = 3347 ± 165, n = 5 mice, n >39 neurons per mouse, linear mixed effects model, p<00001) (**Fig. 1B**).

**Fig 1:**
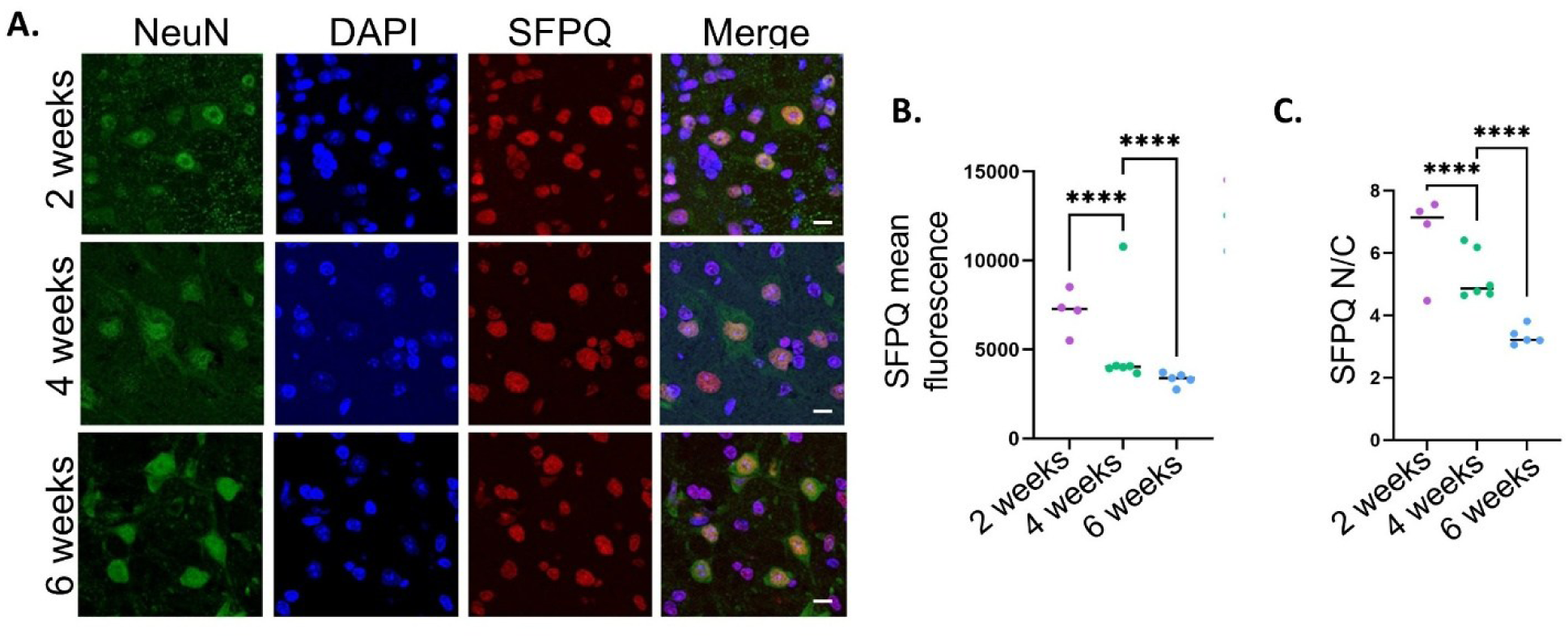
SFPQ expression is reduced and undergoes an early cytoplasmic shift in the TDP-43^rNLS8^ mouse model. **A.** Representative image of mouse spinal cord immunostainined with SFPQ antibody (red), NeuN neuronal marker (green) and DAPI nuclear marker (blue). Scale bar =10 pm. **B.** Whole cell SFPQ expression was significantly reduced between 2 weeks (mean = 7150 + /-622, n = 4 mice, n > 56 neurons per mouse) and 4 weeks (mean = 5094 +/-1139, n = 6 mice, n > 121 neurons per mouse, p <0.0001) and again, between 4 and 6 weeks (mean = 3347 +/-165, n = 5 mice, n >39 neurons per mouse, p<00001). **C.** SFPQ is a predominantly nuclear protein with a mean nuclear / cytoplasmic ratio (N/C) of 6.6 +/-0.7 at 2 weeks post induction. A significant cytoplasmic shift in expression was observed by 4 weeks (mean = 5.3 +/-0.33, p<00001). This shift significantly greater by 6 weeks (mean = 3.3 +/-0.13, p<00001).

The subcellular localisation of SFPQ, quantified as the nuclear-to-cytoplasmic (N/C) ratio, revealed a significant shift towards the cytoplasm between 2 weeks (mean = 6.6 ± 0.7) and 4 weeks post induction (mean = 5.3 ± 0.33; *p* < 0.0001). This redistribution became more pronounced by 6 weeks post induction (mean = 3.3 ± 0.13; linear mixed effects model, *p* < 0.0001), (**Fig. 1C).** Examination of compartment-specific expression of SFPQ at each timepoint demonstrated that this altered ratio was a cumulative result of a reduction in relative nuclear expression (2 week mean = 2.7± 0.11 4 week mean = 2.5 ± 0.08, 6 week mean = 1.8 ± 0.02), and an increase in relative cytoplasmic SFPQ expression (2 week mean = 0.51 ± 0.04 4 week mean = 0.57 ± 0.02, 6 week mean = 0.63 ± 0.03), Supplementary Fig. 1B-C).

These findings are consistent with established *in vitro* studies of TDP-43 LOF^9,19^ and demonstrate that alterations in SFPQ expression and localisation occur rapidly in response to disease-relevant TDP-43 pathology, prior to onset of overt neuronal death at 6 weeks post induction (Supplementary Fig.1). This temporal relationship suggests that SFPQ alterations may actively contribute to neuronal loss, rather than purely representing a late-stage consequence of TDP-43 pathology in this model.

### SFPQ loss of function affects TDP-43 transcript usage and protein expression

Two consistently reported features of SFPQ pathologies are nuclear depletion (LOF) and cytoplasmic accumulation. We first examined the effect of SFPQ LOF on TDP-43 expression at the RNA and protein levels.

### SFPQ LOF is associated with differential TARDBP transcript usage

While TDP-43 loss-of-function (LOF) is known to drive aberrant SFPQ RNA processing^19,21^, whether there is a reciprocal effect of SFPQ LOF on TDP-43 RNA processing remains unknown. To investigate this, HEK293T cells were transfected with siRNA targeting SFPQ, or a scrambled siRNA control (n = 3 replicates). RNA was extracted 48 hours post transfection and short read RNA sequencing (RNAseq) performed.

Principal component analysis demonstrated clear separation between each condition, indicating consistent gene expression changes across biological replicates (Supplementary Figure 2A). SFPQ expression was shown to be significantly reduced by ∼4-fold (log_2_ fold change =-2.099) in the SFPQ siRNA relative to the scramble siRNA treated group, confirming effective knockdown (**Fig. 2A**). Differential gene expression (DGE) analysis identified widespread transcriptional changes following SFPQ LOF, with 43.8% (9666/22090) of tested genes significantly differentially expressed (adjusted p value < 0.05) between SFPQ siRNA and scramble treated groups (Supplementary File 1). When considering genes with an absolute log_2_ fold change >1, 548 (2.5%) and 1531 (6.9%) genes were significantly up and downregulated, respectively. Despite these broad transcriptomic alterations, total *TARDBP* mRNA levels were unchanged between SFPQ knockdown and scrambled control cells (**Fig. 2A**).

**Fig 2:**
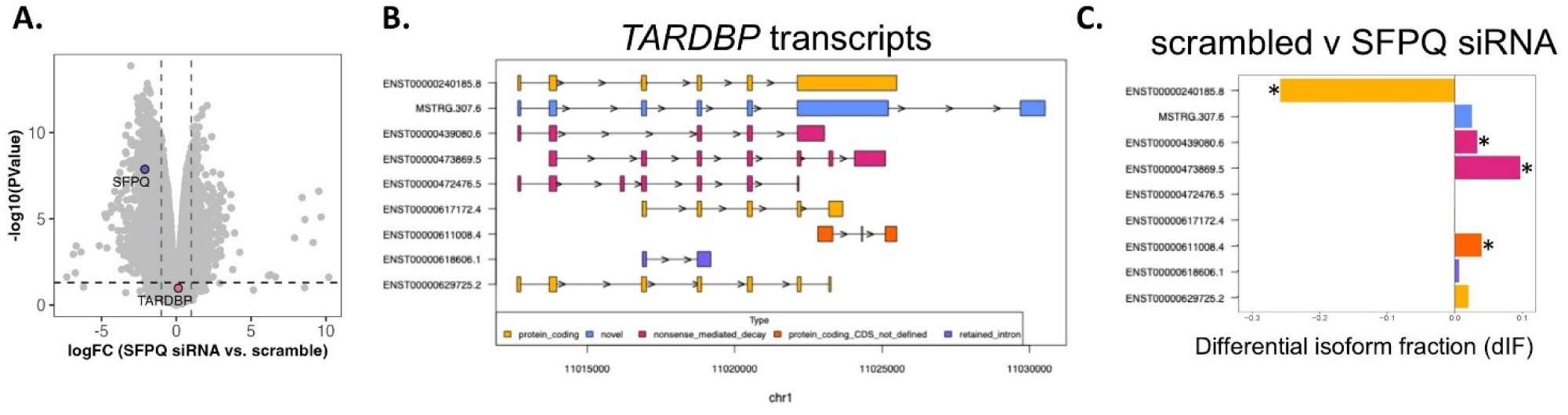
SFPQ loss of function is associated with a shift in TDP-43 RNA isoform usage towards a transcript susceptible to nonsense-mediated decay. **A.** Volcano plot showing gene expression changes in HEK-293T cells treated with SFPQ siRNA relative to scrambled control. *SFPQ* gene expression (blue) is downregulated ∼4-fold indicating successful knockdown. *TARDBP* gene-level expression (red) is not significantly altered. Horizontal dashed line indicates the significance threshold (adjusted p-value < 0.05) and vertical dashed lines indicate two-fold decrease or increase in gene expression. **B.** Visual of **all** *TARDBP* transcripts identified by RNAseq coloured according to GENCODE transcript type annotation or StringTie novel transcript prediction. **C.** Differential isoform fraction of each detected *TARDBP* isoform in HEK-293T cells treated with SFPQ siRNA relative to scrambled control. In SFPQ siRNA treated cells, usage of the canonical *TARDBP* transcript was significantly reduced (dIF--0.259, isoform switch q value = 5.5e-09) while expression of a transcript susceptible to nonsense-mediated decay was significantly increased (dIF = 0.097, isoform switch q value = 3.1e-05).

We next considered whether there were shifts in the abundance of individual *TARDBP* transcript isoforms relative to total gene expression. In addition to the canonical protein-coding transcript, eight *TARDBP* isoforms were detected, including one novel isoform predicted by StringTie **(Fig. 2B**). Differential transcript usage analysis revealed a significant reduction in the use of the canonical *TARDBP* transcript in SFPQ-depleted cells relative to the scramble siRNA control (ENST00000240185.8, 25.9% reduced usage). This was accompanied by a corresponding increase in the usage of an isoform GENCODE annotated as subject to nonsense-mediated decay (NMD; ENST00000473869.5, 9.7% increased usage) (**Fig. 2C**, individual replicates shown in Supplementary Figure 2B).

Given this shift toward an NMD-susceptible isoform, we next asked whether expression of the TDP-43 protein was reduced following SFPQ knockdown.

### SFPQ LOF is associated with reduced TDP-43 protein expression across multiple models

To determine whether SFPQ LOF impacts TDP-43 at the protein level, we first analysed protein lysates collected from HEK293T cells 48 hours post transfection with either SFPQ-targeting or scrambled siRNA (n = 3 per condition). Western blotting confirmed efficient SFPQ knockdown in siRNA-treated cells (**Fig. 3A**; Supplementary Fig. 3A), with SFPQ protein levels reduced by approximately 66% relative to scrambled siRNA controls (0.44 ± 0.05; **Fig. 3B**). This was accompanied by a 36% reduction in TDP-43 protein expression (mean = 0.64 ± 0.04) in SFPQ siRNA treated cells compared to controls (**Fig. 3C**).

**Fig 3:**
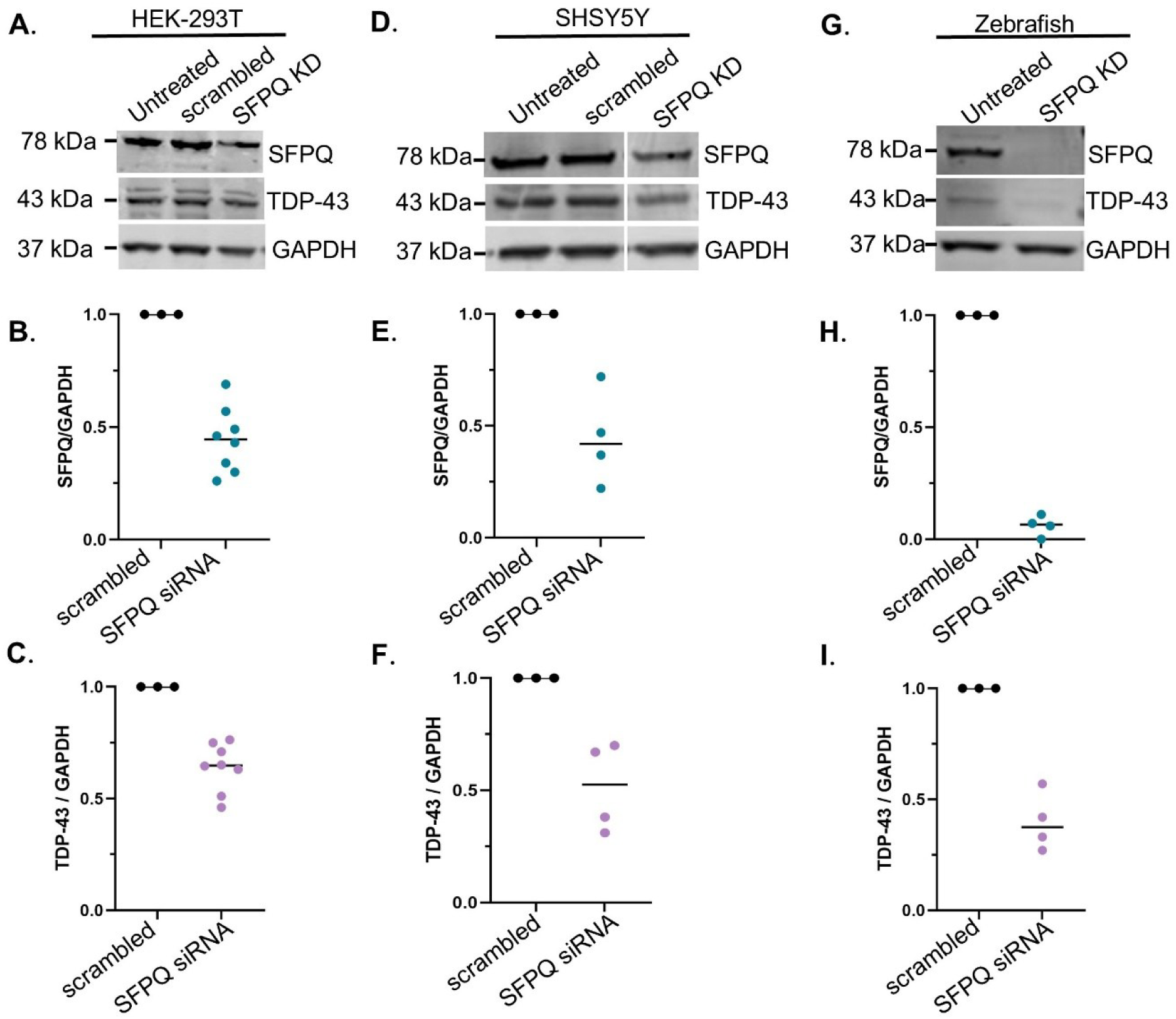
SFPQ loss of function is associated with reduced TDP-43 expression. **A.** Western blot of HEK-293T cells transfected with siRNA targeting SFPQ. **B.** SFPQ knockdown normalised to GADPH loading control was confirmed (mean 0.44 +/-0.14). **C.** This was associated with reduced TDP-43 expression relative to cells treated with scrambled control (mean = 0.64+/-0.11). **D.** Western blot of SHSY5Y cells transfected with siRNA targeting SFPQ. **E.** SFPQ knockdown normalised to GAPDH loading control was confirmed (mean = 0.45 +/-0.11). **F.** This was associated with reduced expression of TDP-43 relative to cells treated with scrambled control (mean 0.52 +/-0.1). **G.** Western blot of zebrafish whole body lysates collected at 2 dpf injected with gRNA targeting SFPQ and cas9 endonuclease (CRISPR-cas9 knockout). **H.** SFPQ knockdown normalised to GAPDH loading control was confirmed (mean = 0.06 +/-0.02 relative to uninjected controls) **I.** This was associated with reduced expression of TDP-43 relative to uninjected controls (mean 0.4 +/-0.07).

To determine whether this finding was conserved in a neuronal-like cellular context, we next transfected SH-SY5Y cells under the same conditions (n = 3-4 per condition; **Fig. 3D**, Supplementary Figure 3B). SFPQ expression was reduced by 55% (mean = 0.45 ± 0.11) relative to scrambled siRNA controls (**Fig. 3E**) and this was associated with a 48% decrease in TDP-43 protein levels (mean 0.52 ± 0.1, **Fig. 3F**).

To establish whether the effect extended beyond cell-based systems and was biologically relevant *in vivo,* we next knocked down SFPQ expression in zebrafish via CRISPR–Cas9 genome editing. Protein lysates were collected at 24 hours post-fertilisation (hpf) and western blotting performed (n = 4 technical replicates; 30 larvae per replicate; **Fig. 3G**, Supplementary Figure 3C). SFPQ protein levels were reduced by ∼94% relative to uninjected controls (mean = 0.06 ± 0.02 **Fig. 3H**), which corresponded with a ∼60% (mean = 0.4 ± 0.07) decrease in TDP-43 expression (**Fig. 3I).**

Therefore, despite differences in the degree of gene suppression achieved by siRNA-mediated knockdown and CRISPR-mediated knockout, both approaches produced concordant effects, providing robust evidence demonstrating that SFPQ LOF reduces TDP-43 protein expression,

### Cytoplasmic accumulation of SFPQ promotes SFPQ phase separation and TDP-43 redistribution

Having examined the effect of SFPQ LOF on TDP-43, we next investigated whether a second hallmark of SFPQ pathology, cytoplasmic accumulation^12–15,17,18,22^, also influences TDP-43. We first characterised the behaviour of SFPQ following its accumulation in the cytoplasm under physiologically relevant conditions by inducing mislocalisation through oxidative stress via sodium arsenite (NaAsO₂). We then assessed the consequences of cytoplasmic SFPQ accumulation for TDP-43 using an SFPQ construct lacking the C-terminal NLS (EGFP–SFPQ^ΔNLS^) in cell lines and zebrafish motor neurons.

### Stress induces SFPQ mislocalisation and accumulation largely independent of stress granules, ubiquitination and TDP-43

NaAsO₂ (50 μM, 20 h) redistributed SFPQ from nucleus to cytoplasm in HEK-293T cells, significantly reducing the SFPQ N/C ratio from 24 ± 2.3 to 14 ± 0.13 (n = 22-23 p = 0.0004, Mann–Whitney) and producing discrete cytoplasmic SFPQ puncta absent in controls (**Fig. 4A, 4B**). The same response was observed in SH-SY5Y cells, where the N/C reduced from 25 ± 2.1 to 1.5 ± 0.22 (n = 20–26; p < 0.0001; **Fig. 4C, 4D**).

**Figure 4:**
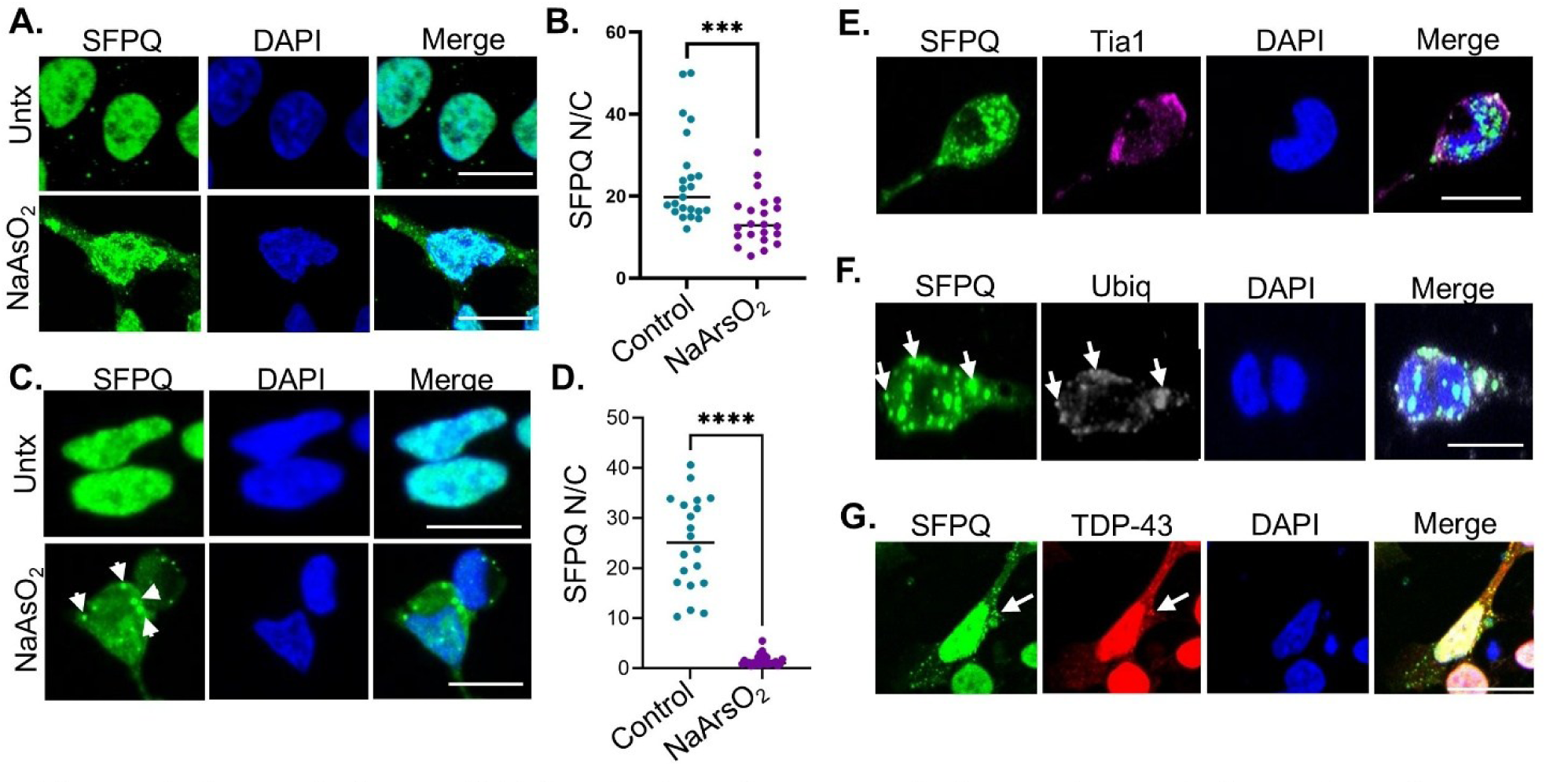
Stress induces SFPQ cytoplasmic accumulation and puncta formation. **A.** Representative images of SFPQ in HEK-293T cells exposed to NaAsO_2_ stress (50 pM 24 hours) demonstrating a cytoplasmic shift of SFPQ and the formation if discrete SFPQ puncta (arrows, Scale bar = 10pm). **B.** Quantification of the nuclear-cytoplasmic ratio of SFPQ demonstrated that the cytoplasmic shift of the protein is statistically significant (untreated mean = 24 +/-2.3, treated mean = 14 +/-1.3, p = 0.0004 n = 22-23). **C.** Representative images of SFPQ in SH-SY5Y cells exposed to NaAsO_2_ stress (50 pM 24 hours). **D.** Quantification of the nuclear-cytoplasmic ratio of SFPQ in SH-SY5Y cells demonstrated that the cytoplasmic shift of the protein is statistically significant (untreated mean = 25 + 2.1 to 1.5 ± 0.22, treated mean = 1.5 + 0.22, p < 0.0001 n = 20-26 cells). **E-G.** Immunofluorescent staining of NaAsO_2_ treated SH-SY5Y cells. Scale bar = 10 pm **E.** Cytoplasmic SFPQ puncta rarely (9% of SFPQ puncta) co-localised with the Tia1 stress granule marker. **F.** Cytoplasmic SFPQ puncta occasionally co-localised with ubiquitin (6% of SFPQ puncta, n = 22 puncta of 367) **G.** Cytoplasmic SFPQ puncta occasionally co-localised with TDP-43 33% of SFPQ positive cytoplasmic puncta (24 of 73 puncta).

To determine whether these puncta corresponded to stress granules, immunofluorescence staining was performed in the SH-SY5Y cells using antibodies targeting the stress granule marker TIA-1 and SFPQ. While robust stress granule formation was evident in NaAsO₂-treated cells (**Fig. 4E**), only 9% of SFPQ puncta co-localised with TIA-1 (37 of 411 of SFPQ positive puncta) within the 20 hour timeframe.

We next investigated whether cytoplasmic SFPQ puncta were associated with protein quality control pathways. Co-staining SFPQ with a pan-ubiquitin antibody demonstrated co-localisation in ∼ 6 % of SFPQ positive puncta (n = 22 puncta of 367). While rare, this does suggest that a small subset of mislocalised SFPQ puncta are targeted for proteasomal degradation within 20 hours^23^.

Finally, we assessed whether SFPQ and TDP-43 co-localise under oxidative stress. Consistent with previous reports, NaArO_2_ treatment induced cytoplasmic redistribution of TDP-43 and the formation of discrete TDP-43 positive puncta^24^. However, TDP-43 was identified in only 33% of SFPQ positive cytoplasmic puncta (24 of 73 puncta) (**Fig. 4G**). These findings suggest that, although both proteins undergo stress-induced mislocalisation and cytoplasmic puncta formation, they do not substantially co-localise at this timepoint.

### Cytoplasmic SFPQ forms dynamic condensates independent of stress

To examine the consequences of SFPQ cytoplasmic accumulation independently of external stress and over an extended timeframe, we next established transgenic zebrafish that expressed human EGFP–SFPQ^ΔNLS^ selectively in motor neurons under the *-3mnx1* promoter^25^. Confocal imaging at 2 days post fertilisation (dpf) confirmed predominantly cytoplasmic localisation of SFPQ^ΔNLS^ in motor neurons of the zebrafish and the formation of discrete SFPQ accumulations (**Fig. 5A**). These accumulations were predominantly cytoplasmic (mean cytoplasmic accumulations per motor neuron = 7 ± 0.39), although nuclear accumulations were also observed (mean nuclear accumulation per motor neuron = 2 ± 0.22) (**Fig. 5B**).

**Figure 5:**
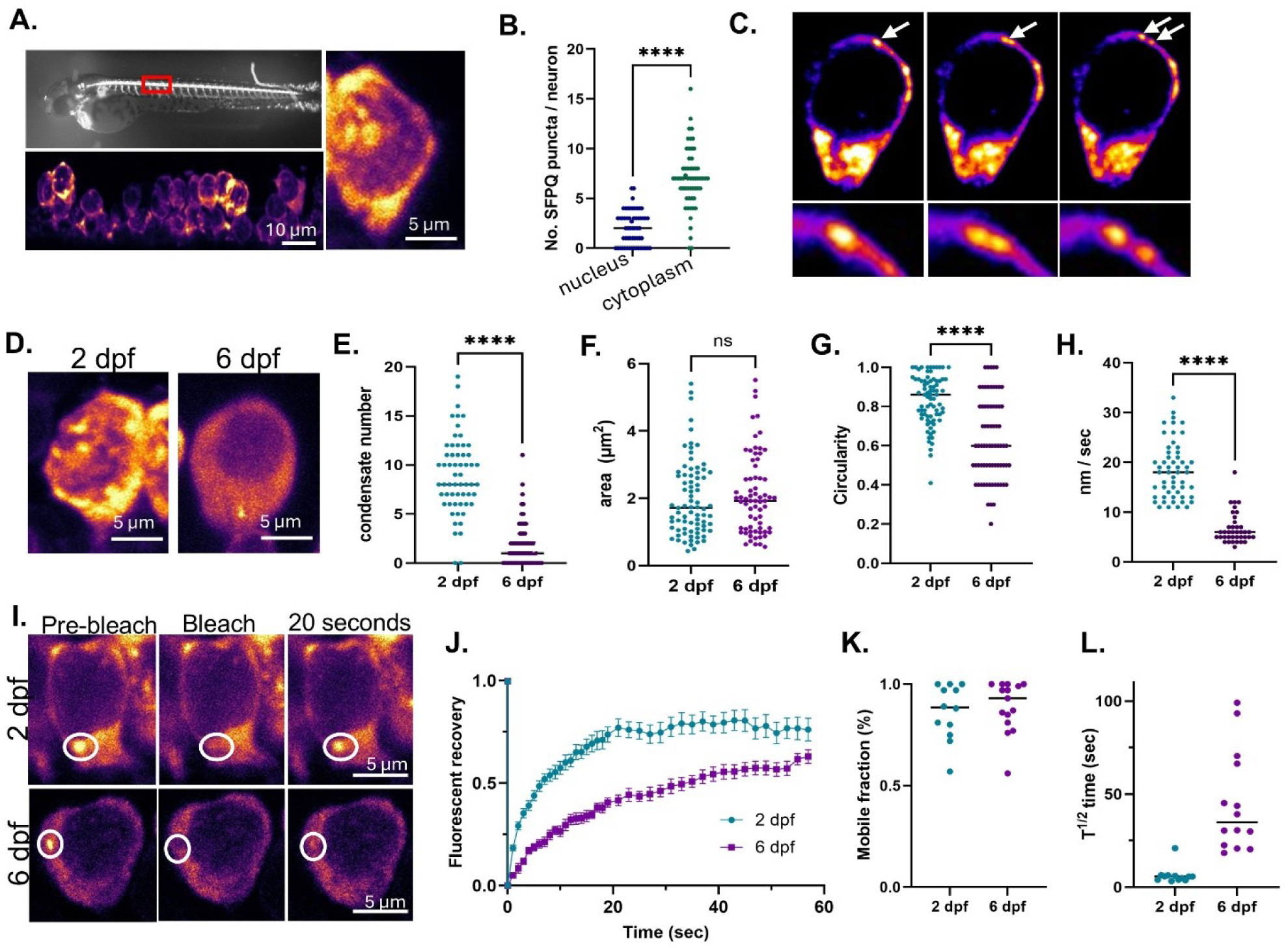
Cytoplasmic SFPQ forms biomolecular condensates that undergo time-dependent changes. **A.** Expression of EGFP-tagged SFPQ^ANLS^ in zebrafish motor neurons at 2 days post fertilisation (dpf) resulted in predominantly cytoplasmic localisation and formation of discrete SFPQ condensates. **B.** SFPQ condensates were observed predominantly in the cytoplasm (mean = 7.0 ± 0.39 condensates per neuron) but were also detected in the nucleus (mean = 2.0 ± 0.22 condensates per neuron). **C.** Time-lapse imaging demonstrated that SFPQ condensates were highly dynamic and underwent fusion and fission events, consistent with biomolecular condensate behaviour. **D - H. SFPQ condensate properties change between 2 and 6 dpf. D.** Representative images at each timepoint. **E.** The number of cytoplasmic SFPQ condensates decreased significantly between 2 dpf (7.0 ± 0.39 condensates, n = 59 neurons) and 6 dpf (1.8 ± 0.25 condensates, *n* = 72 neurons, p<0.0001, Mann-Whitney test). **F.** Average condensate size was unchanged between 2 dpf (2.0 ±0.13 pm^2^, *n* = 79 condensates) and 6 dpf (2.1 ± 0.14 pm^2^, *n* = 69 condensates, p = 0.49 Mann—Whitney test). **G.** Condensate morphology changed over time, with circularity decreasing from 0.83 ± 0.014 at 2 dpf *(n =* 78 condensates) to 0.63 ± 0.024 at 6 dpf *(n =* 72 condensates; *p <* 0.0001, Mann-Whitney test). **H.** Condensate mobility was significantly reduced at 6 dpf (6.7 ± 0.5 nm/s, n = 39 neurons) compared with 2 dpf (18.0 ± 0.67 nm/s, *n* = 54 neurons; *p <* 0.0001, Mann-Whitney test). I - L. **Fluorescence recovery after photobleaching (FRAP) analyses of SFPQ condensates.** I. Representative images of fluorescence recovery after photobleaching (FRAP) analyses at 2 and 6 dpf. **J.** Trace of fluorescent recovery in the 50 seconds post-bleaching. Note At 2 dpf, the high mobility of condensates limited reliable tracking beyond 50s, as structures frequently moved out of the imaging plane. Extended plot of 6 dpf recovery shown in Supplementary fig 4). **K.** The mobile fraction (maximum fluorescence recovery) did not differ between 2 dpf (0.86 ± 0.04, *n* = 12 condensates) and 6 dpf (0.89 ± 0.033, *n* = 16 condensates). L. In contrast, the half-time to fluorescence recovery (f^1^/_2_) increased significantly from 6.4 ± 1.4 s at 2 dpf *(n* = 12) to 45 ± 7.2 s at 6 dpf (n = 16; *p <* 0.0001, Mann-Whitney test).

Time-lapse imaging at 2 dpf demonstrated that the cytoplasmic SFPQ condensates were highly dynamic, moving an average of 18 nm / sec, and fission–fusion events were observed (**Fig. 5C**), consistent with the behaviour of biomolecular condensates. Expression of EGFP–SFPQ^ΔNLS^ in human SY-SY5Y cells similarly induced formation of dynamic SFPQ condensates which underwent fission-fusion events (Supplementary Figure 4B.).

While SFPQ is known to form biomolecular condensates in the nucleus, most notably as a key component of paraspeckles^26^, SFPQ cytoplasmic condensates are yet to be characterised. To address this, we first assessed whether cytoplasmic SFPQ condensates correspond to stress granules by performing co-localisation analysis of EGFP–SFPQ^ΔNLS^ zebrafish injected with mRNA encoding a stress granule marker, G3BP1 or TIA-1, fused to a mCherry fluorophore. Confocal imaging at 2 dpf revealed no detectable co-localisation between SFPQ condensates and G3BP1 (Supplementary Fig 4B.), suggesting that, cytoplasmic SFPQ condensates form independently of canonical stress granules.

### Cytoplasmic SFPQ condensates exhibit progressive changes in morphology and dynamics over time in zebrafish motor neurons

We next characterised the dynamics of the SFPQ cytoplasmic condensates at two timepoints (**Fig 5D**). Condensate number, size, shape, movement and fluorescence recovery after photobleaching (FRAP) were compared at 2 dpf and 6 dpf. A significant reduction in the number of SFPQ condensates in motor neurons was observed between these timepoints suggesting partial clearance or resolution over time. At 2 dpf, mean condensate number per neuron was 7 ± 0.39 (n = 59 neurons, 6 zebrafish) compared to a 6 dpf mean pf 1.8 ± 0.25, (n = 72, 6 zebrafish) (p<0.0001, Mann-Whitney test for nonparametric data) (**Fig. 5E**).

The average size of the condensates remained unchanged between 2 dpf (mean= 0.2 μm^2^

± 0.13, 79 condensates) and 6 dpf (mean =2.1 μm^2^ ± 0.14, n = 69 condensates, Mann-- Whitney test for nonparametric data, p = 0.49, **Fig. 5F**;). However, notable changes in condensate morphology were detected. At 6 dpf, condensates displayed reduced circularity (mean = 0.63 ± 0.024, n = 72) compared to those observed at 2 dpf (mean = 0.83 ± 0.014, n = 78, p <00001, Mann-Whitney test for nonparametric data, **Fig. 5G)**. In line with a reduced phase separation, condensate motility was significantly decreased at the later timepoint with average movement (nm/ second) reducing from a mean = 18 ± 0.67 at 2 dpf (n = 54 neurons over six zebrafish) to mean = 6.7 ± 0.5 at 6 dpf (n = 39 neurons across 7 zebrafish, p <0.0001, Mann-Whitney test for nonparametric data, **Fig. 5H**).

To assess molecular mobility within the condensates, we next compared fluorescence recovery after photobleaching analysis (FRAP) (**Fig. 5I, 5J**). The mobile fraction (percentage of maximum fluorescence recovery) was unaltered between 2 dpf (mean = 0.86 ±0.04, n= 12) and 6 dpf (mean = 0.89 ± 0.033 n = 16) (**Fig. 5K)**. However, the half-time to maximum fluorescence recovery (t½) increased from 6.4 seconds (±1.4, n = 12) at 2 dpf to 45 seconds (± 7.2, n = 16) at 6 dpf (p < 0.0001, Mann-Whitney test for nonparametric data, **Fig. 5L)**.

Collectively, these analyses of SFPQ dynamics indicate that neurons can partially clear cytoplasmic SFPQ condensates over time in motor neurons *in vivo*. However, condensates that persist undergo progressive changes consistent with reduced dynamics and a shift from a liquid-like to a gel-like state as evidenced by decreased circularity, reduced motility, and delayed FRAP recovery.

### SFPQ^ΔNLS^ induced cytoplasmic accumulation is associated with a cytoplasmic shift of TDP-43

Having characterised the progressive maturation of cytoplasmic SFPQ condensates, we next investigated the consequences of SFPQ mislocalisation for TDP-43 behaviour. To enable simultaneous **visualisation** of both proteins *in vivo*, SFPQ^ΔNLS^ transgenic zebrafish were bred with zebrafish expressing mScarlett3-tagged human TDP-43^WT^ in motor neurons. As previously reported, at 2 dpf TDP-43 exhibited predominantly nuclear localisation in neurons negative for SFPQ^ΔNLS^ ^27,28^, (TDP-43 mean N/C = 3.5 ± 0.14). However, neurons that co-expressed TDP-43 and SFPQ^ΔNLS^ were found to have significantly increased cytoplasmic TDP-43, as evidenced by a TDP-43 N/C mean = 2.6 ± 0.05 (linear mixed-effects model, p< 0.0001, n >10 zebrafish fish per condition, > 10 neurons per zebrafish)

(**Fig. 6A**). To determine whether this effect was specific to TDP-43 or reflected a more general disruption of nucleocytoplasmic transport and/or nuclear envelope integrity, SFPQ^ΔNLS^ zebrafish were crossed with a transgenic line that expressed a fluorescent reporter flanked by two NLS driving nuclear expression (NLS-mScarlet3-NLS). At 2 dpf, the mScarlet3 N/C ratio in SFPQ^ΔNLS^ positive neurons (mean = 2.6 ± 0 0.07) was significantly reduced compared to SFPQ^ΔNLS^ negative neurons (mean = 3.6 ± 0.15), indicating a general cytoplasmic shift of nuclear proteins (linear mixed-effects model, p< 0.0001, n >10 zebrafish fish per condition, > 10 neurons per zebrafish) (**Fig. 6B**).

**Figure 6.**
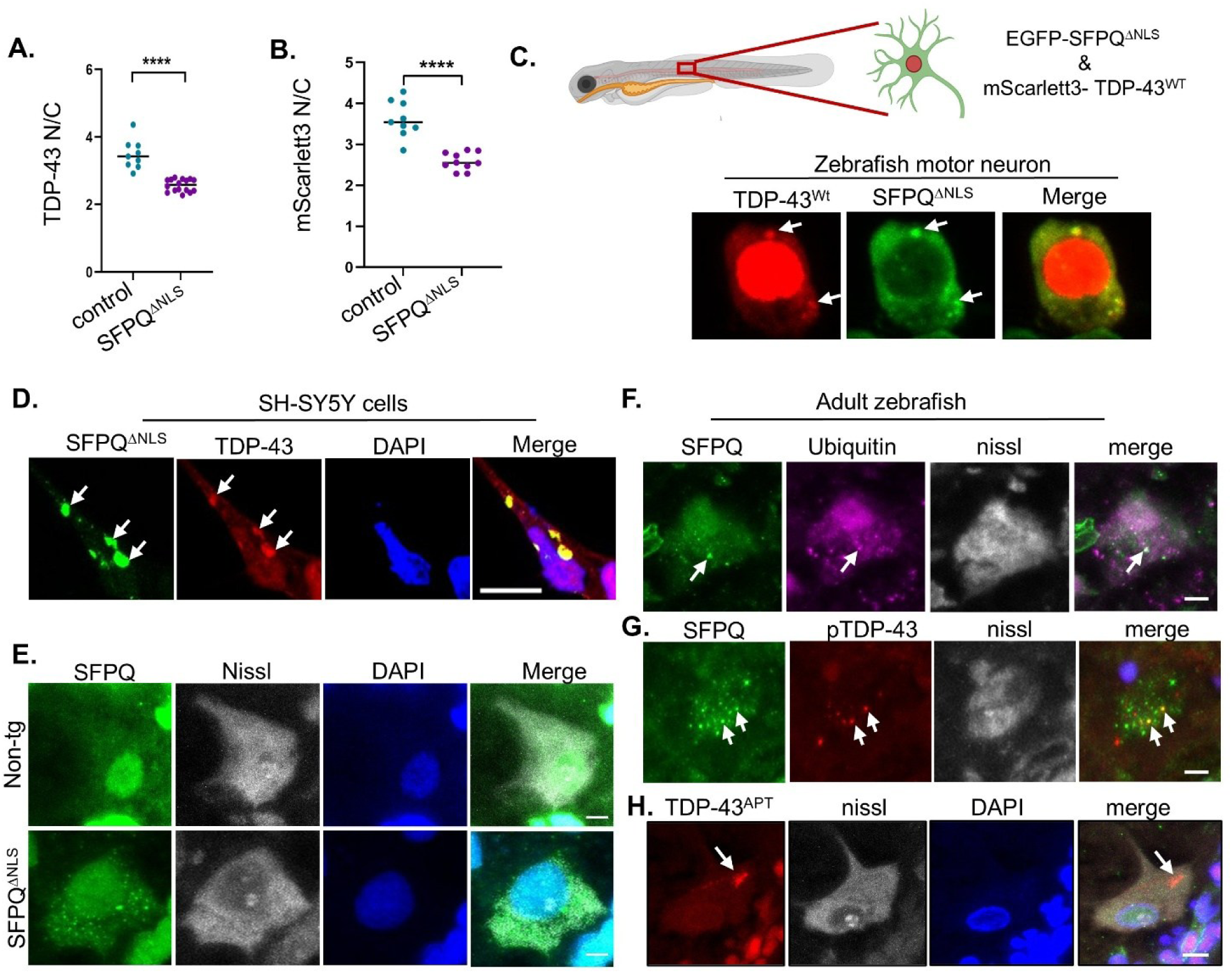
Cytoplasmic SFPQANLS promotes TDP-43 mislocalisation and accumulation of pathological TDP-43 species in motor neurons in vivo. **A.** Quantification of TDP-43 N/C ratio in zebrafish cO-expressing EGFP-tagged SFPQ^ANLS^ and mScarlett3-tagged TDP-43 within motor neurons, significant reduction in the TDP-43 N/C ratio was observed in SFPQ positive neurons (mean = 2.6 ± 0.05) relative to SFPQ“ negative neurons (mean = C = 3.5 ± 0.14, p = p< 0.0001, n = 10 fish, >10 neurons per fish). **B.** Quantification of the N/C ratio of a fluorescent nuclear localisation reporter (NLS-mScarlett3-NLS). Reporter localisation was significantly reduced in SFPQ’ positive neurons (mean *-2.6* ± 0 0.07) relative to SFPQ ^>NLS^ negative neurons (mean = 3.6 ± 0.15. p p< 0.0001, n = 10 fish, >10 neurons per fish). **C.** Approximately 35% of cytoplasmic SFPQ condensates recruited TDP-43 in zebrafish motor neurons (13 of 37 condensates). **D.** Recruitment of endogenous TDP-43 was also observed in SH-SY5Y cells expressing SFPQANLS, with approximately 58% of cytoplasmic SFPQ condensates larger than 1 pm containing TDP-43 (21 of 36 condensates). **E-H.** Immunostaining of adult zebrafish spinal cord sections. Scale bar = 5 pm. **E.** Endogenous SFPQ formed cytoplasmic puncta in SFPQ zebrafish rarely observed in non transgenic zebrafish. **F.** A subset of SFPQ cytoplasmic puncta co-localised with ubiquitin (XX% of puncta). **G.** A subset of SFPQ cytoplasmic puncta co-localised with phosphorylated TDP-43 (pTDP-43) (XX% of puncta). **H.** Staining with a conformation-specific TDP-43 aptamer^designed to detect misfolded TDP-43 species, revealed cytoplasmic accumulations in SFPQ zebrafish neurons (14 of 38 neurons) that were absent from controls.

### Cytoplasmic SFPQ condensates can sequester TDP-43

Confocal microscopy of zebrafish co-expressing EGFP-SFPQ^ΔNLS^ and mScarlett3-TDP-43^WT^ demonstrated recruitment of TDP-43 into cytoplasmic SFPQ condensates in approximately 30.7% of double-transgenic motor neurons (16 of 52) (**Fig. 6C**). To validate this observation in a complementary human cellular model, co-localisation analysis was also performed in human SH-SY5Y cells transfected with SFPQ^ΔNLS^ and immunostained for endogenous TDP-43. No evidence of TDP-43 recruitment within small SFPQ cytoplasmic puncta (< 1 μm^2^) was observed, however larger SFPQ condensates were frequently (65.4% of condensates, 36 of 55) found to contain TDP-43 (**Fig. 6D**).

### Aged SFPQ^ΔNLS^ zebrafish motor neurons develop pathological TDP-43 species

To assess the consequences of cytoplasmic SFPQ accumulation on TDP-43 beyond 6 dpf, SFPQ^ΔNLS^ and non-transgenic control zebrafish were aged to nine months post-fertilisation at which point spinal cord sections were examined. We used a SFPQ antibody that binds to the C-terminal NLS sequence of SFPQ (and therefore does not recognise the SFPQ^ΔNLS^ transgene) to compare endogenous SFPQ. Analysis revealed the presence of endogenous

SFPQ positive cytoplasmic puncta in SFPQ^ΔNLS^ zebrafish that were largely absent from controls (**Fig. 6E**).

Ubiquitin staining revealed limited association with the endogenous SFPQ puncta, (∼7,5%, 27 of 360 SFPQ puncta) (**Fig. 6F**, Supplementary Figure 5A), suggesting that while some structures may engage protein quality control pathways, widespread ubiquitination is not a defining feature at this stage in this model.

Notably, phosphorylated TDP-43 was detected in a subset of SFPQ-positive puncta (**Fig. 6G**, Supplementary Figure 5B) indicating that persistent SFPQ assemblies can, in some cases, recruit pathological forms of TDP-43 over time. However, co-localisation events were infrequent, with only 7.5% of SFPQ puncta (12 of 160 SFPQ-positive puncta) also positive for pTDP-43. Consequently, the majority of SFPQ condensates remained pTDP-43-negative. Importantly, no pTDP-43 skein-like inclusions were observed.

To further investigate possible pathological TDP-43 alterations in the SFPQ^ΔNLS^ neurons, spinal cord sections from 9-month-old zebrafish were labelled with a TDP-43 aptamer probe that recognizes conformationally altered TDP-43 preceding overt misfolding and aggregation^29,30^. Discrete cytoplasmic TDP-43 accumulations were observed in neurons of SFPQ^ΔNLS^ zebrafish (n = 38 neurons examined, n = 14 with TDP-43 positive accumulations, **Fig. 6H**, Supplementary Figure 5B). Similar accumulations were not observed in non-transgenic zebrafish of the same age.

## Discussion

TDP-43 pathology is a defining feature of multiple neurodegenerative diseases, yet the upstream mechanisms that initiate and propagate TDP-43 dysfunction remain incompletely understood. Here, we demonstrated that SFPQ perturbation is both an early consequence of TDP-43 pathology and a driver of TDP-43 dysfunction. Specifically, we showed that SFPQ LOF altered *TARDBP* transcript processing, promoting production of an isoform predicted to undergo NMD (**Fig.2**). Consistent with this, we showed that SFPQ LOF reduces TDP-43 protein abundance (**Fig.3**). In parallel, our data demonstrated that cytoplasmic accumulation of SFPQ promotes TDP-43 mislocalisation and recruitment into pathological assemblies in motor neurons *in vivo* (**Fig.6**). Collectively, these findings indicate that SFPQ dysfunction influences both the abundance and subcellular distribution of TDP-43.

When considered alongside previous studies showing that TDP-43 LOF drives SFPQ dysregulation through altered *SFPQ* polyadenlyation^19,21^, our findings support a model in which SFPQ and TDP-43 form a reciprocal regulatory axis. We propose that disruption of either protein initiates a self-reinforcing feed-forward cycle, whereby TDP-43 dysfunction drives SFPQ pathology, which then feeds back to exacerbate TDP-43 dysfunction, ultimately amplifying disease progression.

### SFPQ LOF and cytoplasmic accumulation are early consequences of TDP-43 pathology

Nuclear depletion and cytoplasmic mislocalisation of SFPQ have been reported in patient studies^12,13,15,17,18^ and experimental models of ALS, FTD and AD^12–15^, implicating SFPQ dysfunction as a common feature of neurodegeneration. However, it remains unclear whether these abnormalities represent an early pathogenic event or arise secondary to neurodegeneration. Using the inducible TDP-43-rNLS mouse model^20^, we observed SFPQ dysregulation relatively early in the disease course. In this model, expression of cytoplasmic TDP-43 and concomitant depletion of endogenous nuclear TDP-43 is associated with a well characterised pathological sequence where presymptomatic neuromuscular junction defects emerge at 2 weeks post-induction, motor impairment at 4 weeks, motor neuron loss at 6 weeks, and end-stage disease by 8–10 weeks^20^. In this study, reduced neuronal SFPQ expression and a redistribution of SFPQ from the nucleus to the cytoplasm were evident by 4 weeks post-induction, coinciding with the onset of motor dysfunction and preceding significant motor neuron loss. A probable mechanism for the SFPQ perturbations comes from previous transcriptomic analyses showing that TDP-43 LOF induced alternative polyadenylation of *SFPQ*, which generated a NMD-sensitive transcript and, in transcripts that escape degradation, an SFPQ protein lacking a nuclear localisation signal^9,19^.

While our data establish SFPQ perturbation as an early consequence of TDP-43 pathology, it does not resolve whether dysfunction of one protein precedes the other or whether both arise in response to common upstream pathogenic stressors. Defining the temporal sequence of these events will require longitudinal studies with sufficient resolution to capture the earliest changes in both proteins.

### SFPQ Loss of function altered *TARDBP* / TDP-43 expression

SFPQ has multiple RNA synthesis and RNA processing functions. These include regulation of transcription initiation, transcript elongation (especially of long genes > 100kb)^34,35^, RNA splicing^31–33^ and regulation of alternative polyadenylation site usage ^34,35^. Consequently, SFPQ LOF has widespread effects on the transcriptome. With respect to *TARDBP*, we found that total *TARDBP* RNA abundance was unchanged following SFPQ knockdown, however expression of the canonical protein-coding transcript was reduced and accompanied by a reciprocal increase in alternative isoforms predicted to undergo NMD (**Fig.2**). These findings suggest that SFPQ is not required for *TARDBP* transcription but instead plays an important role in *TARDBP* transcript processing.

The mechanism underlying this processing defect remains to be determined. SFPQ may directly regulate *TARDBP* pre-mRNA processing through interactions with the transcript itself, or alternatively the observed splicing changes may arise indirectly through broader alterations to RNA processing pathways following SFPQ depletion. Regardless of the underlying mechanism, our data demonstrated that SFPQ LOF altered *TARDBP* transcript usage, establishing dysregulated *TARDBP* processing as a significant consequence of SFPQ dysfunction.

The shift to non-productive *TARDBP* transcripts provides the most plausible explanation for the robust reduction in TDP-43 protein abundance that we observed across multiple SFPQ knockdown model system (**Fig. 3).** Tight regulation of TDP-43 abundance is essential for cellular homeostasis - even modest perturbations can disrupt hundreds of RNA processing events implicated in neurodegeneration^2^. Consequently, multiple regulatory mechanisms have evolved to maintain TDP-43 within a narrow physiological range. These include an autoregulatory feedback loop^36,37^, regulation by other RNA-binding proteins and splicing factors^38^, and epigenetic control^39^. Our findings identify SFPQ as an additional component of this essential regulatory network.

### Oxidative stress as a potential initiator of SFPQ nuclear LOF and cytoplasmic accumulation

One plausible upstream contributor to SFPQ mislocalisation, and thereby nuclear LOF and cytoplasmic accumulation, is oxidative stress - an early and persistent feature of neurodegenerative disease^40,41^. Consistent with this hypothesis, NaArO_2_ exposure induced cytoplasmic redistribution of SFPQ and the formation of SFPQ-positive puncta that were largely distinct from canonical stress granules. This susceptibility of SFPQ to oxidative stress may arise from its molecular composition. SFPQ contains redox-sensitive cysteine residues within its intrinsically disordered low-complexity domains and is enriched in methionine residues, both of which are susceptible to oxidative modification^42,43^. Such modifications could alter SFPQ localisation, intermolecular interactions and liquid-liquid phase separation behaviour.

Notably, oxidative stress is a well-established driver of TDP-43 mislocalisation^24,44^, raising the possibility that cellular stress represents a common upstream event that initiates both SFPQ and TDP-43 pathological changes.

### SFPQ mislocalisation promoted condensate formation, disruption of nuclear–cytoplasmic homeostasis and TDP-43 transition to pathological states

Our data indicated that cytoplasmic accumulation of SFPQ has the capacity to initiate multiple pathogenic processes relevant to TDP-43 proteinopathies. We demonstrated that, when present at sufficient concentration, SFPQ undergoes liquid–liquid phase separation within the cytoplasm. SFPQ is well recognised as a component of nuclear condensates and is essential for the formation and maintenance of paraspeckles, dynamic nuclear ribonucleoprotein bodies involved in regulating gene expression, RNA processing, and cellular stress responses^45^. To the best of our knowledge, however, this is the first demonstration of SFPQ condensate formation in the cytoplasm outside of its previously described association with stress granules^43^. These findings suggest that cytoplasmic mislocalisation of SFPQ not only depletes its normal nuclear functions but may also confer novel, potentially pathogenic effects.

One potential consequence of SFPQ cytoplasmic condensate formation is disruption of nuclear–cytoplasmic protein homeostasis. A major finding of this study was that cytoplasmic SFPQ accumulation drove redistribution of TDP-43 from the nucleus to the cytoplasm of motor neurons *in vivo* (**Fig.6**). Given that cytoplasmic TDP-43 mislocalisation is a defining pathological feature of TDP-43 proteinopathies^4^, this observation is particularly significant.

Importantly, however, the effect was not restricted to TDP-43. SFPQ cytoplasmic accumulation also induced redistribution of a nuclear-targeted fluorescent reporter (NLS-mScarlet3-NLS) to the cytoplasm, suggesting a broader disturbance of nuclear–cytoplasmic compartmentalisation rather than a TDP-43-specific mechanism.

Although cytoplasmic SFPQ accumulation has not previously been linked to defects in nucleocytoplasmic transport, studies of TDP-43 provide potential insight into the mechanisms underlying this observation. Cytoplasmic TDP-43 aggregates sequester nucleoporins and other components of the nuclear transport machinery. However, disruption of this system is also evident in the absence of overt aggregation^46^. Changes in the phase separation behaviour of TDP-43 has been shown to cause sequestration of importin-α and the nucleoporin protein Nup62, causing a progressive impairment nuclear import^44^. Whether SFPQ condensates similarly recruit components of the nuclear transport machinery and thereby impair nucleocytoplasmic trafficking remains an important question for future investigation.

Beyond this hypothesised effect on protein localisation, our data suggested that cytoplasmic SFPQ condensates may contribute to the formation of pathological protein aggregates. We observed that cytoplasmic SFPQ condensates displayed progressive changes in their biophysical properties, transitioning from highly mobile structures with rapid fluorescence recovery to assemblies that were less spherical, less mobile and slower to recover following photobleaching. These features are consistent with reduced molecular exchange and decreased solubility (**Fig.5**). Current models propose that pathological TDP-43 inclusions form through the maturation of physiological condensates into gel-like and ultimately insoluble aggregates. Under cellular stress, TDP-43 undergoes intra-condensate de-mixing, promoting local unfolding, intermolecular cross-linking, and a liquid-to-gel transition. Further structural destabilization and loss of RNA binding enhance aberrant hydrophobic and electrostatic interactions, leading to the formation of increasingly insoluble fibrillar aggregates^47^. The progressive reduction in solubility of the SFPQ condensates observed in this study may suggest a similar process occurs upon prolonged SFPQ accumulation in the cytoplasm. Supporting this concept, SFPQ aggregates, both TDP-43 positive and negative, have been identified in patients affected by ALS^16^ and Alzheimer’s disease^15^.

Finally, TDP-43 was recruited into SFPQ condensates in both zebrafish motor neurons and human neuron-like cells, suggesting that SFPQ condensates may act (at least moderately) as sites that concentrate and sequester TDP-43, potentially increasing its susceptibility to pathological conversion. Supporting this hypothesis, aged SFPQ^ΔNLS^ zebrafish developed multiple hallmarks of TDP-43 pathology. These included the presence of occasional phosphorylated TDP-43-positive puncta and cytoplasmic accumulations of misfolded TDP-43 detected using a conformation-specific TDP-43 aptamer^29,30^. Importantly, these pathological TDP-43 species were absent from control animals. Together, these findings suggest that persistent cytoplasmic SFPQ accumulation creates an intracellular environment that promotes TDP-43 mislocalisation, sequestration and pathological conversion, rather than simply becoming incorporated into pre-existing aggregates.

### Outstanding Questions and Future Directions

Critical questions remain regarding the relationship between TDP-43 and SFPQ in the context of neurodegenerative diseases. For example, it is not yet clear whether one pathology commonly emerges before the other, or whether SFPQ alteration represent an under-recognised accelerator of disease progression. Addressing these questions will be essential for therapeutic concepts, as effective intervention strategies may depend on targeting these contributors sequentially or with different intensities to interrupt this pathogenic cycle.

While our data support a role for SFPQ in regulating *TARDBP* transcript processing, TDP-43 protein expression, and localisation, the precise molecular mechanisms driving these effects remain to be defined. In addition, although we demonstrate that SFPQ dysregulation can drive key features of TDP-43 pathology, the extent to which this contributes to neuronal dysfunction and degeneration requires further investigation. In addition, while zebrafish models provide valuable insights into dynamic cellular processes, additional characterisation in mammalian systems will be important to establish translational relevance. In particular, integrating temporal analyses of SFPQ and TDP-43 localisation in other human cell and mouse models may further refine our understanding of disease progression.

## Summary

In summary, this study identifies SFPQ as a central regulator of TDP-43 homeostasis and provides mechanistic insight into how its dysregulation can drive hallmark features of TDP-43 proteinopathies. By linking SFPQ loss of function, cytoplasmic condensation, and progressive TDP-43 pathology, our findings support a model in which SFPQ operates within a bidirectional, feed-forward loop that amplifies neurodegenerative processes. Targeting this axis may therefore represent a novel strategy for modifying disease progression in ALS and related disorders.

## Methods

### Histopathological analysis of TDP-43-rNLS mouse spinal cord tissue

#### Mouse generation and *tissue preparation*

Mice used in this study were bred and maintained at The University of Queensland under approved animal ethics protocols (QBI/131/18 and 2021/AE000200). All experimental procedures were conducted in accordance with the Australian Code for the Care and Use of Animals for Scientific Purposes and were approved under animal ethics approval 2022/AE000578.

TDP-43 rNLS8 double transgenic mice^20^ and single transgenic littermate controls on a pure C57Bl/6JAusb background were produced as previously described^48^ through breeding of transgenic mice expressing the tetracycline transactivator (tTA) under the control of the human neurofilament heavy chain (*NEFH*) promoter with mice carrying a tetracycline-responsive (*tetO*) human TDP-43 transgene harbouring a defective nuclear localisation signal (hTDP-43ΔNLS). The resulting bigenic offspring (rNLS mice) exhibit doxycycline-regulated expression of cytoplasmic hTDP-43ΔNLS in neurons of the brain and spinal cord. Mice were maintained on Dox-containing chow until transgene induction by doxycylcline withdrawal at approximately 8 weeks of age.

At the designated experimental timepoints (2, 4 and 6 weeks off doxycycline), mice were euthanised and spinal cords extracted. Tissues were fixed in 10% neutral-buffered formalin, processed routinely, and embedded in paraffin wax. Paraffin-embedded spinal cords were sectioned at 6 μm thickness using a rotary microtome and mounted on glass slides.

Given the absence of reported sex-dependent differences in the TDP-43-rNLS mouse model, both male and female mice were included. Spinal cord sections were collected at 2, 4, and 6 weeks after doxycycline withdrawal. Experimental cohorts consisted of non-transgenic mice at 2 weeks (n = 5 females), 4 weeks (n = 4 females, n = 1 male), and 6 weeks (n = 4 females, n = 1 male) off doxycycline, and TDP-43-rNLS mice at 2 weeks (n = 2 females, n = 2 males) and 6 weeks (n = 3 females, n = 2 males) off doxycycline.

#### Immunofluorescent staining

Formalin-fixed paraffin-embedded tissue sections were deparaffinised by incubation at 60°C for 35 min, followed by rehydration through sequential washes in xylene (3 × 10 min), 100% ethanol (3 × 5 min), 70% ethanol (5 min), and Milli-Q water (2 × 3 min). Antigen retrieval was performed in pH 6.0 citrate buffer (Sigma-Aldrich) at 96°C for 20 min.

Sections were washed three times in 0.1% Triton X-100 in phosphate-buffered saline (PBS-T) for 5 min each at room temperature. A hydrophobic barrier was drawn around each section using a PAP pen (Sigma-Aldrich), and sections were blocked in 5% normal goat serum (NGS) in PBS-T for 1 h at room temperature in a humidified chamber.

Primary antibodies diluted in 5% NGS/PBS-T were applied to sections and incubated overnight at 4°C 1:200 rabbit SFPQ (ab38148), 1:200 mouse NeuN (ab104224). Sections were then washed in PBS-T before incubation with secondary antibodies (1:250 goat anti-rabbit Alexafluor 555 and goat anti-mouse Alexafluor 488) for 1 h at room temperature.

Followign PBS washes, sections were mounted with Fluoroshield containing DAPI (Sigma-Aldrich).

#### Imaging and analysis

Slides were imaged at 40^ magnification using the Olympus Slideview VS200. Image analysis was performed using QuPath (v0.6.0) with the Cellpose extension (v4.0.6) employing the default Cyto3 model. The areas of the ventral horn was manually delineated, then automated batch analysis conducted with viable motor neurons defined as cells containing a single DAPI-positive nucleus within a NeuN-defined cytoplasmic boundary.

For each neuron, QuPath extracted cell area (μm²), and whole-cell, nuclear, and cytoplasmic SFPQ fluorescence intensity. Raw output data were imported into RStudio (v2025.05.1+513) for downstream analysis.

### In vitro assays

#### siRNA SFPQ knockdown in HEK293T cells

HEK293T and SH-SY5Y cells were maintained in Dulbecco’s Modified Eagle Medium (DMEM; Sigma Aldrich) supplemented with 10% fetal bovine serum (Sigma Aldrich) at 37 °C, 5% CO_2_ and 95% humidity. DsiRNA targeting SFPQ (5’ – GAAGGUGUUUUCUUACUGACGACAA, 3’ – CACUUCCACAAAAGAAUGACUGCUGUU) and a universal scrambled control were purchased from Integrated DNA Technologies (IDT, Coralville, IA, USA).

Cells were seeded in 6 well dishes for a confluency at transfection ∼ 30%. Transfections were performed using Lipofectamine RNAiMAX transfection reagent (Theromofisher) in opti-MEM media (Life Technologies) as per manufacturer’s recommendation, with a final siRNA concentration of 10 µM.

#### RNA extraction and sequencing

At 48 hours post transfection, HEK-293T were washed in ice-cold PBS, then Buffer RLT (RNeasy kit, QIAGEN) with 1:100 β-mercaptoethanol (β-ME) was added to each well. Cells were manually detached with a cell scraper, collected and vortexed. RNA was extracted from these samples using the RNeasy kit (QIAGEN) as per manufacturer’s instructions.

Samples were stored at –80 °C until samples from three replicates were collected.

Sample quality control, library preparation and RNA sequencing was performed by the Australian Genome Research Facility/Foundry (AGRF, Sydney Australia). Total RNA quality was assessed using an Agilent TapeStation system, with all samples found to have high integrity (RNA Integrity Number equivalent [RINe] = 10). Sequencing libraries were prepared using the Illumina Total RNA Library Preparation kit with Ribo-Zero Plus.

Sequencing was performed on an Illumina NovaSeq X Plus platform to a minimum depth of approximately 100 million reads per sample (range: 141 – 174 million 150 bp paired-end reads).

#### RNAseq analysis

Raw FASTQ files were processed using Trimmomatic v0.39^49^ to remove low-quality reads and adaptor sequencers. Quality checks were performed using FastQC v0.11.9 (http://www.bioinformatics.babraham.ac.uk/projects/fastqc/), multiQC v1.12^50^ and Picard CollectRnaSeqMetrics tool (http://broadinstitute.github.io/picard).

Trimmed FASTQ files were aligned to release 48 of the GENCODE human reference genome using STAR v2.7.10b with --twopassMode Basic parameter specified ^51^. Gene and transcript counts were generated using RSEM v1.3.3^52^ and were read into R using tximport v1.38.2^53^. To consider de novo transcripts, StringTie v3.0.3^54^ (excluding - e option) was used to assemble transcripts from STAR genome-aligned BAM files.

StringTie was then run in merge mode to generate a combined transcriptome annotation containing both known and de novo transcripts. Transcripts were re-assembled using the de novo transcriptome as reference and StringTie-eb options, generating read coverage tables. All following analyses were performed in R v4.5.2.

Following filtering of low count genes (<10 counts in <3 samples), gene counts were normalised using a variance stabilizing transformation^55^. Principal component analysis was performed on the resulting expression matrix using prcomp and samples were visualised by 2D plotting of principal components to check for outliers.

Differential gene expression analysis was performed between SFPQ siRNA and scramble groups using edgeR v4.8.2^56^. Genes with an adjusted p-value (FDR) < 0.05 were considered differentially expressed.

To identify transcripts with differential usage in SFPQ siRNA versus scramble treated cells, IsoformSwitchAnalyzeR v2.10.0^57^, was applied to StringTie-derived transcript quantification using the DEXSeq^58^ implementation of the isoform switch test. Significant isoform switches were defined as FDR corrected p-value (q-value) < 0.05 and ≥ 10% change in isoform usage. *TARDBP* transcript structures and isoform fractions were visualised with a custom R script and ggplot2^59^ respectively.

#### Protein extraction

For protein lysate collection from cell lines, HEK293T or SH-SY5Y cells were seeded into 6 well plates and transfected with siRNA as described. 48 hours post transfection, wells were washed with ice-cold phosphate buffered saline (PBS). PBS was replaced with RIPA buffer (20 mm Tris-HCl, pH 7.4, 1% Triton X-100, 150 mm NaCl, 1 mM EDTA) with complete protease inhibitor cocktail, Roche Applied Sciences) and cells physically detached with a cell scraper. Lysates were intermittently vortexed for 30 minutes at 4°C and then probe sonicated. The homogenised sample was centrifuged for 20 minutes at 13,000 rpm at 4°C and the supernatant collected for analysis.

#### Western blotting

Protein concentration was determined using the Pierce BCA Protein Assay Kit and western blotting performed as previously described. Briefly, 30 µg protein lysate was loaded into a 4-15% SDS-PAGE gel (Biorad) and electrophoresed for 5 minutes at 50V followed by 40 minutes at 180V. Samples were transferred to a nitrocellulose membrane via wet transfer. Membranes were blocked in 0.5% Tween 20, 3% BSA in PBS for 1 hour at room temperature, followed by overnight incubation in primary antibody at 4°C. Primary antibodies used were rabbit anti-SFPQ (1:800), rabbit anti-TDP-43 (1:10000 Proteintech, 10782-2-AP), and rabbit anti-GAPDH (1:15000, Proteintech, 60004-1-Ig). The following day, membranes were washed in 0.5% Tween 20 and then incubated for one hour at room temperature in IRDye 800CW donkey anti-rabbit IgG and 680LT donkey anti-mouse IgG, 1:15000 (LCR-926-32212 and LCR-926-68023 respectively, LI-COR Biosciences).

Membranes were imaged on the Odyssey CLx imaging system and protein bands quantified with the Image Studio Lite software (LI-COR).

#### Application of oxidative stress

HEK293T and SHSY5Y cells were seeded onto poly-L-lysine treated coverslips in a 24-well plate. At confluency of ∼ 50%, NaArsO_2_ was applied at 50 μM for 20 hours followed by PBS washes and fixation.

#### Transfection of SFPQ^ΔNLS^ constructs

The human SFPQ gene sequence (**NM_005066.3**) with the C-terminal nuclear localisation sequence (NLS) deleted was synthesised by Genscript (previously described ^60,61^) and cloned into a pcDNA3.1 vector with a N-terminal EGFP fluorescent tag. HEK293T and SH-SY5Y cells were seeded into a 24 well plate with coverslips and transfected using Lipofectamine LTX reagent with PLUS (Thermofisher Scientific). as per manufacturer’s instructions.

#### Immunofluorescent staining of cell lines

At 24 hours post application of stress or 48 hours post DNA transfections, culture medium was aspirated and cells were concurrently fixed and permeabilised in 0.2% triton X and 4% PFA in PBS for 10 minutes at room temperature. Following PBS washes, cells were blocked in 5% NGS for 1 hour at room temperature. Primary antibodies used were: rabbit anti-SFPQ (1:800, Sigma-Aldrich ab38148), rabbit anti-TDP-43 (1:1000; 10782-2-AP), mouse anti-phosphorylated Ser409/410 TDP-43 (1:5000, TIP-PTD-M01, Cosmo Bio), mouse anti-ubiquitin (1:150, MAB1510, Merk Millipore,MA, US) and goat anti-TIA-1 (1:500 Santa Cruz sc-1751). Following overnight incubation at 4^○^C, cells were washed in PBS and incubated in species-appropriate secondary antibody (AlexaFluor488 or AlexaFluor555, 1:250) for one hour at room temperature. Coverslips were mounted in Fluoroshield with DAPI (Sigma-Aldrich) and imaged on a ZEISS LSM 880 inverted confocal laser-scanning microscope.

### Zebrafish assays

#### Zebrafish ethics and husbandry

Zebrafish used in this study were bred and maintained under established conditions^62^ and all husbandry and experimental procedures were performed in compliance with the Animal Ethics Committee, Macquarie University (NSW, Australia) and the Animal Ethics Committee, University of Sydney (NSW, Australia). All fish used were AB/Tübingen wild type background.

Larvae were maintained at 28°C in E3 embryo medium (5 mM NaCl, 0.17 mM KCl, 0.33 mM CaCl_2_, and 0.33 mM MgSO_4_ buffered to pH 7.3) until six days post fertilisation (dpf). At 6 dpf, larvae were moved onto system water, temperature 28°C, pH 7.4 and housed in 14 hour light-10 hour dark cycle with twice daily artemia and pellet feeds.

#### CRISPR-cas9 SPFQ knockdown in zebrafish

For SFPQ knockdown in zebrafish, three Alt-R CRISPR-Cas9 sgRNA targeting SFPQ were custom designed and purchased from IDT. Sequences shown in **Table 1**.

|  |  |
| --- | --- |
| SFPQ Alt-R CRISPR-Cas9<br>sgRNA # 1 | mA*mC*mG* rArCrG rGrGrC rUrUrG rUrUrC rUrGrU rGrCrG<br>rUrUrU rUrArG rArGrC rUrArG rArArA rUrArG rCrArA rGrUr<br>U rArArA rArUrA rArGrG rCrUrA rGrUrC rCrGrU rUrArU rCrA<br>rA rCrUrU rGrArA rArArA rGrUrG rGrCrA rCrCrG rArGrU rCr<br>GrG rUrGrC mU*mU*mU* rU |
| SFPQ Alt-R CRISPR-Cas9<br>sgRNA # 2 | mG*mC*mG* rArCrG rGrUrG rGrGrA rGrUrC rUrGrC rUrGrG<br>rUrUrU rUrArG rArGrC rUrArG rArArA rUrArG rCrArA rGrUr<br>U rArArA rArUrA rArGrG rCrUrA rGrUrC rCrGrU rUrArU rCrA<br>rA rCrUrU rGrArA rArArA rGrUrG rGrCrA rCrCrG rArGrU rCr<br>GrG rUrGrC mU*mU*mU* rU |
| SFPQ Alt-R CRISPR-Cas9<br>sgRNA # 3 | mC*mG*mC* rArArC rCrUrC rUrGrC rCrArA rGrGrC rArCrG<br>rUrUrU rUrArG rArGrC rUrArG rArArA rUrArG rCrArA rGrUrU<br>rArArA rArUrA rArGrG rCrUrA rGrUrC rCrGrU rUrArU rCrAr<br>A rCrUrU rGrArA rArArA rGrUrG rGrCrA rCrCrG rArGrU rCr<br>GrG rUrGrC mU*mU*mU* rU |

All three sgRNAs (1,000 pg total, 333 pg each) were combined with TrueCut Cas9 protein v 2 (800pg, Thermo Fisher Scientific, Waltham, MA, USA) and incubated at 37^○^C to allow complex formation as previously described ^63^. Microinjection was performed by injecting 0.5-1 nL of the mixture zebrafish eggs at the single cell stage of development.

#### Protein lysate collection

Embryos were manually dechorinated at one dpf, euthanised in 50 mg/L tricaine (MS222) and rinsed in PBS before being deyolked by submersion in deyolking buffer (55 mm NaCl, 1.8 mm KCl, 1.25 mm NaHCO_3_) with agitation, followed by multiple washes in ice cold PBS. Embryos were then homogenised in RIPA buffer (20 mm Tris–HCl, pH 7.4, 1% Triton X-100, 150 mm NaCl, complete protease inhibitor cocktail, Roche Applied Sciences) for 2 minutes, probe sonicated and centrifuged 20 min at 13,000 rpm at 4 °C and the supernatant collected for analysis. Western blotting was performed as described for *in vitro* analysis using rabbit anti-SFPQ (1:800), rabbit anti-TDP-43 (1:10000 Proteintech, 10782-2-AP), and rabbit anti-GAPDH (1:15000, Proteintech, 60004-1-Ig).

#### Generation of DNA Constructs for transgenic zebrafish development

DNA constructs for expression in zebrafish were generated using the Multisite Gateway^®^ Three Fragment Vector Construction^64^. To generate the SFPQ and TDP-43 constructs, The pTol2pA2 destination vector ^64^ was recombined with a p5E-*3mnx1* promotor^25^, a middle entry vector (pME_EGFP or pME-mScarlet3)^25^ and a 3’ entry vector (p3E^64^). The SFPQ^ΔNLS^ sequence was synthesised and cloned into the p3E vector by Genscript (Piscataway, NJ, USA). The p3E:TDP-43^WT^ vector which was generated by subcloning a *Bam*HI–*Spe*1-flanked gene string encoding HsaTDP-43 (GeneArt, Thermo Fisher Scientific, Regensburg, Germany) into the *Bam*HI–*Spe*1 sites of p3E-MCS as previously described ^65^. To generate the NLS-mScarlet3-NLS reporter protein, the pTol2pA2 destination vector ^64^ was recombined with a p5E:*elalv3* promotor^25^, a pME: NLS-mscarlet3-NLS vector, in which the NLS-mscarlet3-NLS sequences was synthesised and cloned into the pME vector by Genscript,, and a p3E-polyA vector^64^.

DNA constructs were microinjected into eggs of wild type zebrafish at 20 ng/µl in combination with transposase mRNA (30 ng/µL) and phenol red dye. Embryos were screened for mosaic expression at ∼48 hpf on a M165FC fluorescent stereomicroscope (Leica).

#### G3BP1 mRNA synthesis for stress granule analysis in zebrafish

The zebrafish (*Danio rerio*) *G3BP1 and TIA-1* coding sequences (ZFIN: ZDB-GENE-030131-7452, ZDB-GENE-030131-1506) were synthesised by GeneArt (Thermo Fisher Scientific, Waltham, MA, USA) as an N-terminal fusion with mCherry. The mCherry-tagged fragment was subcloned into the pCS2+ expression vector^66^.

For mRNA synthesis, the pCS2+-mCherry-G3BP1 and pCS2+-mCherry-TIA-1 constructs were linearised with NotI (New England Biolabs, Ipswich, MA, USA) and purified using a QIAquick Gel Extraction Kit (Qiagen, Hilden, Germany). Capped mRNA was generated from the linearised template by *in vitro* transcription using the mMESSAGE mMACHINE SP6 Transcription Kit (Thermo Fisher Scientific), according to the manufacturer’s instructions.

Transcribed RNA was purified by lithium chloride precipitation and resuspended in nuclease-free water for downstream applications. RNA concentration and purity were measured using a QIAxpert spectrophotometer (QIAGEN, Hilden, Germany) and RNA integrity validated via gel electrophoresis.

Purfied mRNA was diluted to 100ng/ul in dH2O and phenol red dye and injected into – *3mnx1*:EGFP-SFPQ^ΔNLS^ eggs at the single cell stage of development at a bolus size of ∼1nL. Embryos were screened for expression at 48 hpf on a M165FC fluorescent stereomicroscope (Leica).

#### Live imaging of zebrafish embryos

For live imaging studies, zebrafish were anaesthetised in 0.01% (w/v) tricaine methansulfonate (MS-222, Sigma) and mounted on their side in 1% low melting point agarose (Thermofisher). Once the agarose set, zebrafish were submerged in E3 embryo medium with tricaine and imaged on Leica Stellaris 5 confocal microscope with a 25X water immersion lens. All analysis was replicated in zebrafish from a minimum of three clutches and only morphologically normal zebrafish were included in the study.

Analysis of SFPQ condensate number, size and motility and TDP-43 localisation was performed in Fiji (Image J) ^67^ using methods previously detailed ^68^.

For Fluorescent recovery after photobleaching (FRAP) analysis, imaging performed on a SP5 confocal microscope using a Leica 40’/0.80 HCX APO L U-V-I water immersion lens and the FRAP wizard of Leica AF software as previously described ^27^. Briefly, the brightest focal plane of a single condensate was selected and the region of interest / bleach area of 20×20 pixels defined. Photobleaching was performed using a 405nm argon laser at 60% intensity for 3 frames (1.293s each frame). To assess recovery, images were taken every 1 second (20 repeats, then every 2 seconds (66 repeats) then every 5 seconds (26 repeats). To account for fluorescence bleaching during imaging, two additional ROIs were selected - a non-fluorescent background region and a representative section of the whole cell.

Fluorescence intensity of all three regions of interest at each timepoint was determined using a custom Fiji (Image J) ^67^ macro and raw results uploaded into EasyFRAP The fluorescent intensities were corrected for overall bleaching and normalized to the pre-bleach intensity and a two-component exponential equation was confirmed to be the best fit to quantify the recovery curves (https://easyfrap.vmnet.upatras.gr/) ^69^. Only images with a gap ratio (total fluorescence remaining after bleaching) >7 and a bleach depth >5.5 were analysed. Fluorescent recovery was full scale normalized, fitted using a double term exponential equation as previously reported ^69^ and plotted in GraphPad Prism with standard error to the mean (SEM) displayed.

#### Immunofluorescent and TDP-43 aptamer staining of adult zebrafish

Adult *-3mnx1*:EGFP-SFPQ^ΔNLS^ zebrafish were euthanised in tricaine on ice for 30 minutes and sectioned into thirds for fixation in 4% PFA for 24 hours at 4^○^C. Following PBS washes, tissue was dehydrated in 15% 6 hours, then 30% sucrose overnight at 4 ^○^C, then mounted in OCT (Optimal Cutting Temperature) and stored at-80 ^○^C. Tissues were sectioned to 50 µm using a Leica CM1950 cryostat and placed in PBS immediately prior to staining.

For immunofluorescent staining, sections were permeabilised in 0.1% Triton X for 10 minutes, then blocked in 10% NGS for 2 hours at room temperature. Primary antibodies used were rabbit anti-SFPQ (1:800, abcam ab38148), rabbit anti-native TDP-43 (1:1000; 10782-2-AP), mouse anti-TDP-43 phosphorylated Ser409/410 (1:5000, TIP-PTD-M01, Cosmo Bio) or mouse ant-ubiquitin (1:250, MAB1510). Incubation was performed overnight at 4°C. After washing, sections were incubated with species-appropriate secondary antibodies (AlexaFluor 488 or AlexaFluor 555, 1:250) for 2 hours at room temperature.

For aptamer-based detection of TDP-43, sections underwent permeabilisation and blocking as described above before incubation with TDP-43^APT^ (1.2 μg/mL) ^29,30^ diluted in blocking buffer for 4 h at room temperature in the dark. Sections were subsequently washed three times in PBS containing 0.1% Triton X-100 (PBS-T) for 5 min each.

Following antibody or aptamer staining, neuronal cell bodies were labelled with NeuroTrace™ 640/660 fluorescent Nissl stain (Thermo Fisher Scientific) diluted 1:150 in PBS for 30 min at room temperature. Sections were washed once in PBS-T for 15 min, followed by three washes in PBS for 10 min each. Sections were mounted using DAKO mounting medium containing DAPI, coverslipped, sealed with clear nail polish, and cured flat in the dark prior to imaging. Images were acquired using a ZEISS LSM 880 inverted laser-scanning confocal microscope.

## Statistical analysis

Statistical analysis was performed and graphs generated using GraphPad Prism 9 (9.2.0). Shapiro-Wilk normality test or the D’Agostino-Pearson normality test was used to assess normality distribution of data for each assay.

For normally distributed data, two-tailed t-tests were used for pairwise comparisons and one-way ANOVAs and Holm-Sidak’s multiple comparison for datasets with >2 groups. For data with non-Gaussian distribution, Kruskal-Wallis test with multiple comparisons was used.

For analyses of TDP-43-rNLS mouse tissues and SFPQ nuclear-to-cytoplasmic (N/C) ratios in zebrafish neurons, where measurements were obtained from large numbers of cells across multiple biological replicates, differences between treatment groups were assessed using a linear mixed-effects model. Treatment was included as a fixed effect and biological replicate as a random effect to account for within-replicate variability and the non-independence of individual cellular measurements.

For all statistical tests significance was taken as *p < 0.05, **p<0.01, ***p < 0.001. Unless otherwise indicated, data values are presented as the mean ± standard error of the mean (SEM).

## Acknowledgements

We thank the Macquarie University Animal Services (MARS) staff for expert animal care, specifically Jason Martin-Powell and Cheryl Song for their daily animal support. We also thank Kelly Foskett, Tyler Chapman and Emily Don for their help in generating molecular constructs.

The graphical abstract was were created with BioRender.com.

Further information and requests for resources and reagents should be directed to the lead contacts Marco Morsch or Alison Hogan.

