## Supplementary figures for "SFPQ dysregulation promotes TDP-43 pathology through a pathogenic feedback loop"

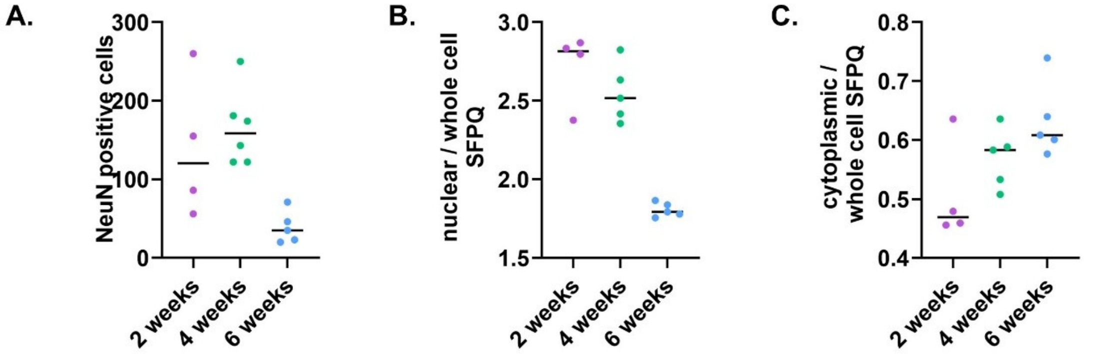


**Supplementary Figure 1: A,** Motor neuron counts in the ventral horn of TDP-43-rNLS8 mice at 2, 4 and 6 weeks. No reduction was observed between 2 weeks post induction (mean NeuN positive cells per section per mouse = 139 +/- 45) and 4 weeks post induction mean NeuN positive cells per section per mouse = 165 +/- 20). At the 6 week timepoint, mean NeuN positive cells per section per mouse had reduced to 39 +/- 9.2) . **B** Fluorescent intensity of SFPQ in neuronal nuclei relative to whole cell SFPQ fluorescent intensity progressively reduced from 2 weeks (2.7 +/- 0.11) to 4 weeks (2.5 +/- 0.083) and to 6 weeks (1.8 +/- 0.02). **C.** Fluorescent intensity of SFPQ in neuronal cytoplasm relative to whole cell SFPQ fluorescent intensity progressively reduced from 2 weeks (0.51 +/- 0.043) to 4 weeks (0.57 +/- 0.022) and to 6 weeks (0.63 +/- 0.028). Collectively, these results indicate that neuronal loss is evident at 6 weeks in our cohort and the observed reduction in SFPQ nuclear cytoplasmic ration is a consequence of a shift of the protein from the nuclear compartment to the cytoplasm.

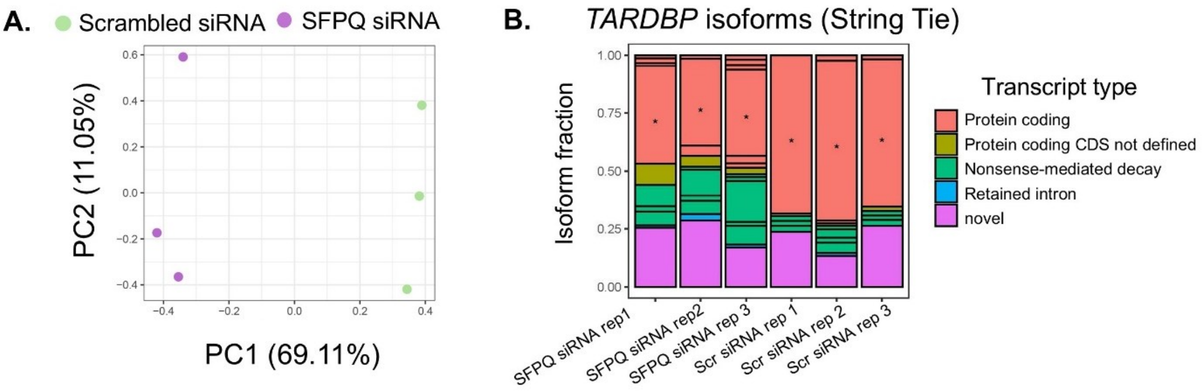


**Supplementary Figure 2: RNAseq analysis of HEK293T cells transfected with SFPQ or scrambled siRNA. A.** Principal component analysis demonstrated clear separation between conditions indicating consistent gene expression changes across biological replicates. **B.** A shift in *TARDBP* isoform usage was consistently observed across replicates. The three SFPQ siRNA treated samples show a significant reduction in the canonical protein coding isoform (indicated by *) and an increased usage of isoforms targeted for nonsense-mediated decay (shown in green), relative to scramble treated controls.

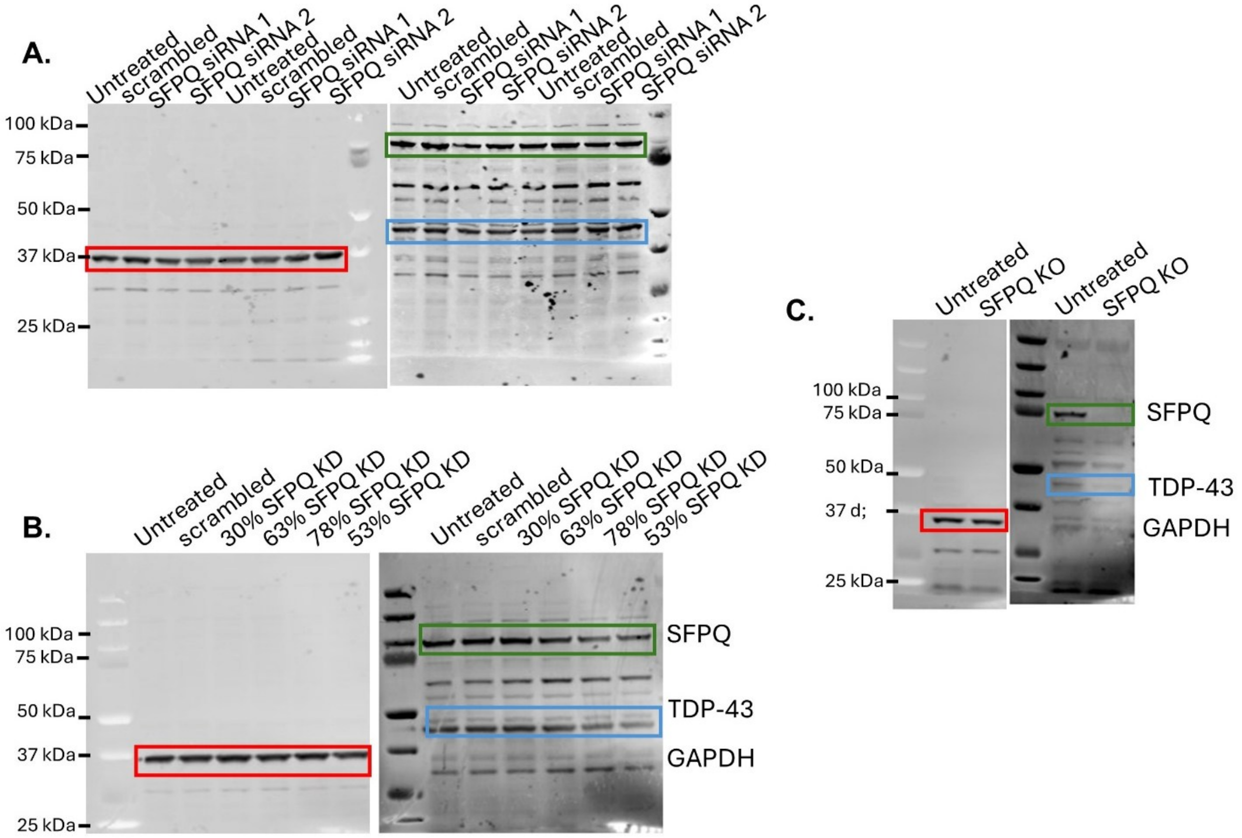


**Supp Fig 3**: Uncropped western blots from **A.** HEK-293T cells transfected with siRNA targeting SFPQ **B.** SH-SY5Y cells transfected with siRNA targeting SFPQ **C.** Zebrafish larvae in which SFPQ was knocked out through CRISPR-cas9. In all blots, GAPDH is indicated by the red box, SFPQ by the green box and TDP-43 by the blue box.

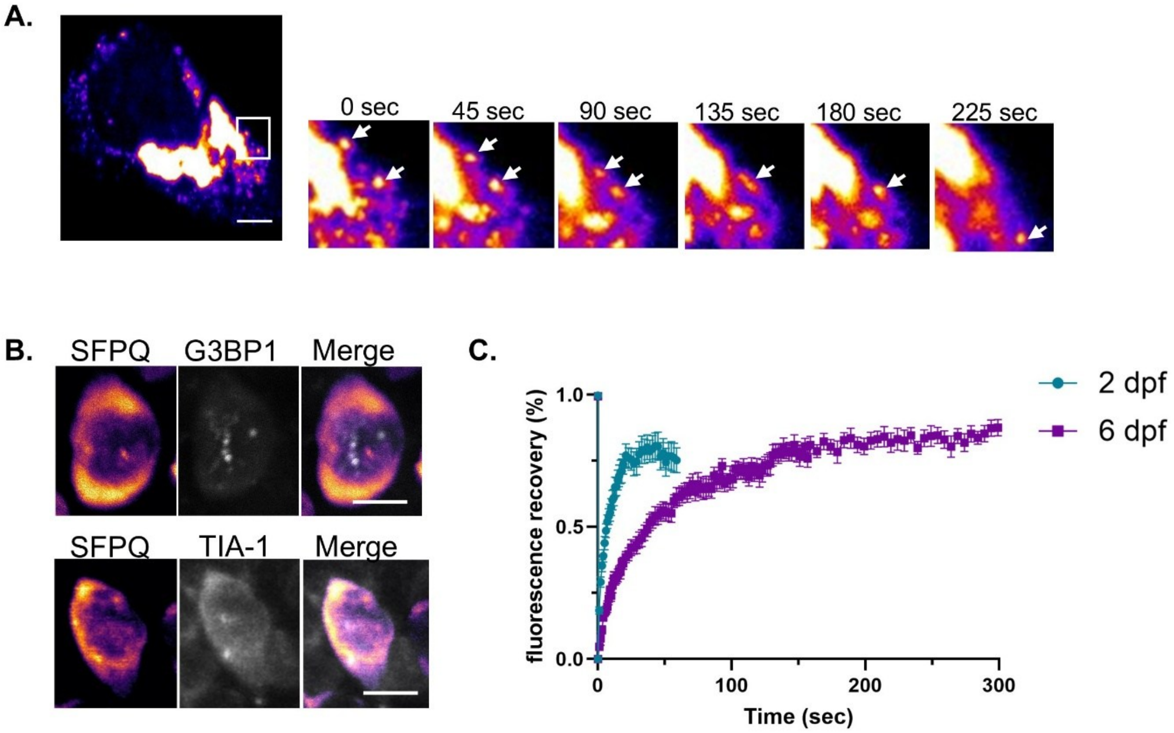


**Supp Fig 4**: **A SH-SY5Y cells transfected with EGFP-tagged SFPQ$^{\Delta NLS}$.** Upon its accumulation in the cytoplasm, SFPQ forms bimolecular condensates *in vitro* which are dynamic and undergo fission-fusion events. **B.** *-3mnx1*:EGFP-SFPQ$^{\Delta NLS}$ zebrafish injected with mRNA encoding mCherry-G3BP1 or m-Cherry-TIA-1. No overlap between SFPQ puncta and either stress granule marker was observed. **C.** Extended FRAP analysis of SFPQ condensates in SFPQ$^{\Delta NLS}$ zebrafish motor neurons. Note that accurate tracking of fluorescent recovery beyond 50 seconds was precluded at 2 dpf by the highly dynamic nature of the condensates.

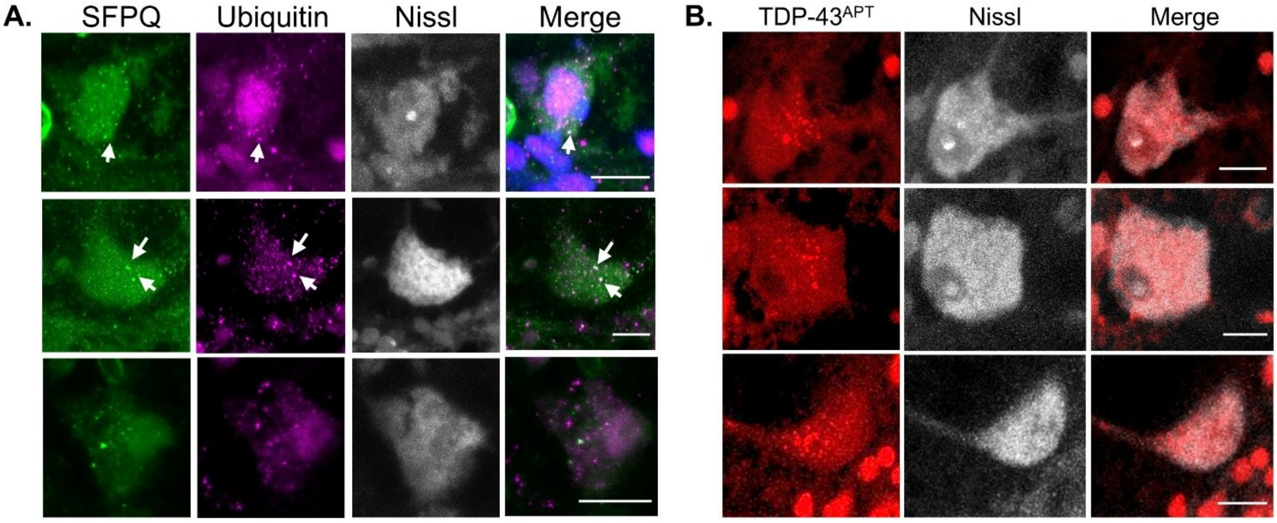


**Supp Fig 5: Additional images of neurons of aged zebrafish expressing SFPQ$^{\Delta NLS}$ .** A. Neurons immunostained for endogenous SFPQ and ubuqiitin (scale bar = 10 μm. **B.** Additional pTDP-43 images. **E.** Additional TDP-43 aptamer images (scale bar = 10 μm. Additional ubiquitin images.
